# Metagenomic strain-resolved DNA modification patterns link extrachromosomal genetic elements to host strains

**DOI:** 10.64898/2026.03.27.714056

**Authors:** Shuai Wang, Allison K. Guitor, Luis E. Valentin-Alvarado, Rebecca E. Garner, Pengfan Zhang, Ming Yan, Ling-Dong Shi, Marie C. Schoelmerich, Holly M. Steininger, Daniel M. Portik, Siyuan Zhang, Jeremy E. Wilkinson, Susan Lynch, Michael J. Morowitz, Matthias Hess, Spencer Diamond, Jillian F. Banfield, Rohan Sachdeva

## Abstract

DNA modification is central to microbial defense against extrachromosomal genetic elements (ECEs), consequently ECEs tend to adopt their host’s modification patterns. Shared ECE–host modification patterns enable linking ECEs to their hosts, but modification detection tools are designed for single genomes and are ineffective at metagenome scale. Here, we present MODIFI, software for detecting DNA modifications in metagenomes. MODIFI assumes that each *k*-mer in a metagenome is mostly unmodified and calculates background signal levels for that *k*-mer from PacBio HiFi reads, eliminating the need for matched control experiments. MODIFI ECE–host linkages were validated using >1,000 isolate and mock microbiome datasets. Illustrating the approach, we identified 315 strain-resolved, non-redundant ECE–host linkages in environmental and human metagenomes. In infant gut microbiomes, a chromosomal inversion in *Enterococcus faecalis* alters host and associated plasmid methylation motifs simultaneously. Overall, MODIFI solves a major bottleneck in DNA modification analysis and provides a foundational tool for understanding microbial epigenomics.

## Main

Linking extrachromosomal genetic elements (ECEs) to their microbial hosts is essential for understanding microbial evolution and adaptation via acquisition of genes from ECEs^1,2^. Current ECE–host linkage strategies are limited. For example, culture-based methods are restricted to the cultivable minority, while sequence-composition^3,4^ and machine-learning^5^ approaches often lack specificity. Although CRISPR-spacer matching^6,7^, proximity ligation (Hi-C), and single-cell sequencing offer higher confidence^8,9^, the former is restricted to taxa with active immune memories, while the latter two remain prohibitively expensive and labor-intensive for large-scale metagenomic studies. Consequently, a scalable, culture-independent method for high-resolution linkage remains critical for an enhanced understanding of ECEs and their biological role.

DNA modification plays a vital role in regulating gene expression and prokaryote defense against ECEs^10–12^. In prokaryotes, restriction modification (RM) systems serve as a primary immune barrier^13^, where methyltransferases (MTases) modify specific motifs to distinguish self from foreign DNA and restriction endonucleases (REases) cleave the foreign DNA. Therefore, ECEs tend to adopt modification patterns of their hosts, and the shared modification patterns can be used for ECE–host linkage^14,15^.

Long-read sequencing technologies offer a powerful approach for the detection of DNA modifications^10,16–18^. PacBio HiFi sequencing, characterized by high base accuracy (∼99.95%)^19^, allows direct detection of modified nucleotides by monitoring polymerase kinetics during the sequencing of the DNA template^16^. However, interpreting polymerase kinetic signatures requires distinguishing true modifications from background variation driven by local sequence context (∼10 bp)^20,21^. Existing methods often rely on whole-genome amplification to generate modification-free controls, which is labor-intensive and costly, or utilize machine-learning models that may not generalize to novel lineages^14,16,22,23^. Furthermore, these tools are primarily optimized for single genomes and lack the scalability required for complex metagenomic datasets^22^. While methylation has been applied to binning and host linkage, no scalable framework exists for high-throughput, whole-metagenome analysis^14,15,24,25^. Additionally, whether methylation can resolve linkages at the strain level, or be applied broadly across diverse phyla and habitats, remains unclear.

Here, we present MODIFI (**MOD**ification un**IFI**er), a software package for strain-resolved DNA modification detection and ECE–host linkage from PacBio metagenomic sequences (Figure 1). MODIFI estimates baseline polymerase kinetics directly from the reads, eliminating the need for matched control experiments and mitigating technical variability across sequencing runs. To systematically link ECEs to their hosts, we developed a linkage score that accounts for the uniqueness of shared modification motifs that are present within a microbial community. By providing efficient, metagenome-scale epigenomic profiling, MODIFI enables precise detection of strain-level epigenetic variation and ECE–host linkage directly from long-read sequences.

**Figure 1.**
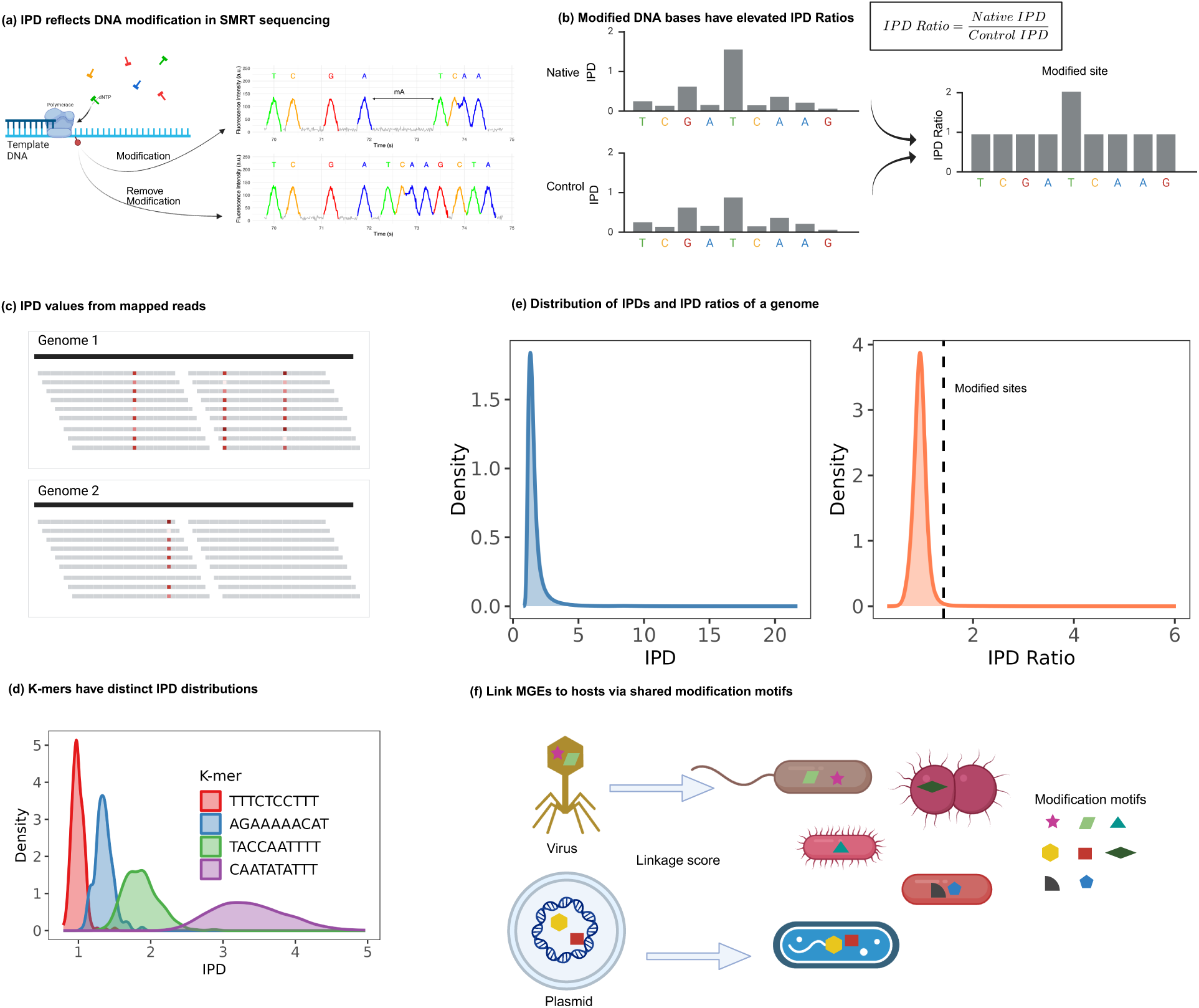
Framework for DNA modification detection in metagenomics. **(a)** Schematic of the PacBio SMRT sequencing process for modification detection; modified DNA bases exhibit increased IPD values. **(b)** Comparison of native DNA IPDs, matched control DNA IPDs, and IPD ratios within a sequence fragment. Elevated IPD ratios specifically highlight the modified bases. **(c)** Visualization of IPD values across reads alignment to reference genomes, with red indicating elevated IPD values. **(d)** IPD distributions for four distinct *k*-mers identified within the same metagenomic dataset. **(e)** Distribution of IPDs and IPD ratios for a specific genome; dashed vertical lines indicate the thresholds used for modification calling. **(f)** Schematic of ECE–host linkage estimation. Host assignments are determined by shared DNA modification motifs between ECEs and candidate genomes.

## Results

### Overview of MODIFI

In PacBio sequencing, modified bases typically exhibit elevated inter-pulse duration (IPD) values (Figure 1a). However, because IPD is influenced by the local DNA sequence context (*k*-mer), an unmodified control baseline is required to calculate the IPD ratio, a normalized metric where elevated values specifically indicate modification (Figure 1b). The MODIFI pipeline begins by aligning reads to a metagenome reference to extract IPD values for each genomic position (Figure 1c). The protocol analyzes both the primary reads (subreads) and consensus HiFi reads. The IPD distribution is distinct for each *k*-mer (Figure 1d). MODIFI establishes an unmodified baseline by assuming that most instances of a *k*-mer at different loci in the metagenome reference are unmodified. Positions with significantly elevated IPD ratio are identified as modified sites (Figure 1e). The associated modification motif is identified *de novo* by comparison of regions flanking the modified sites. Finally, to identify ECE hosts, we calculate a linkage score for each potential ECE– host pair, weighting the associations by the uniqueness of shared methylation motifs within the metagenomic dataset (Figure 1f).

Validation using native and control data from *E. coli* C227 demonstrated that MODIFI accurately detects base modifications across various coverage thresholds in both individual isolate genomes and pooled simulated metagenomes (Supplementary Note S1; Figure S1a, b). In mock microbiome (mixed isolate) benchmarks, MODIFI achieved accurate modification motif detection and maintained high detection sensitivity even at low sequencing depths (Supplementary Note S1; Figure S1c, d; Table S1). Furthermore, MODIFI is highly efficient, processing diverse real-world metagenomes with a mean wall-clock time of 4.5 hours (Supplementary Note S2; Figure S2-S4). Overall, MODIFI provides high accuracy in base modification and motif detection, with high computational efficiency.

### DNA modification is common across habitats and phyla

To assess modification in isolate genomes, we individually analyzed 1,420 pure bacterial isolate genomes (≥10× coverage) representing 153 species across seven phyla, alongside 17 isolates lacking species-level assignments (Figure 2a; Supplementary Table S2). In addition, genomes of two archaeal isolates, *Methanosphaera* sp. and *Natrinema pallidum* were investigated. We found that 96.62% (1,372) of bacterial isolates and both archaeal genomes contained modification motifs. To evaluate modification prevalence in metagenomes from diverse ecosystems, we analyzed 59 PacBio HiFi datasets from human and animal microbiomes, as well as environmental sources such as soil, ocean water, and sugarcane (Supplementary Figure S3, S5; Table S3, S4). From these datasets, we recovered 1,315 high-quality metagenome assembled genomes (MAGs) (≥10× depth, ≥50% completeness, ≤5% contamination) spanning 49 phyla, comprising 1,125 bacterial and 190 archaeal MAGs (Figure 2b, c). Similar to the findings for isolates, 89.13% (1,172/1,315) of the MAGs possessed detectable modification motifs. Analysis of the combined isolate and metagenome datasets reveal that a substantial fraction (93.02%; 2,546/2,737) of genomes contain detectable modification motifs.

**Figure 2.**
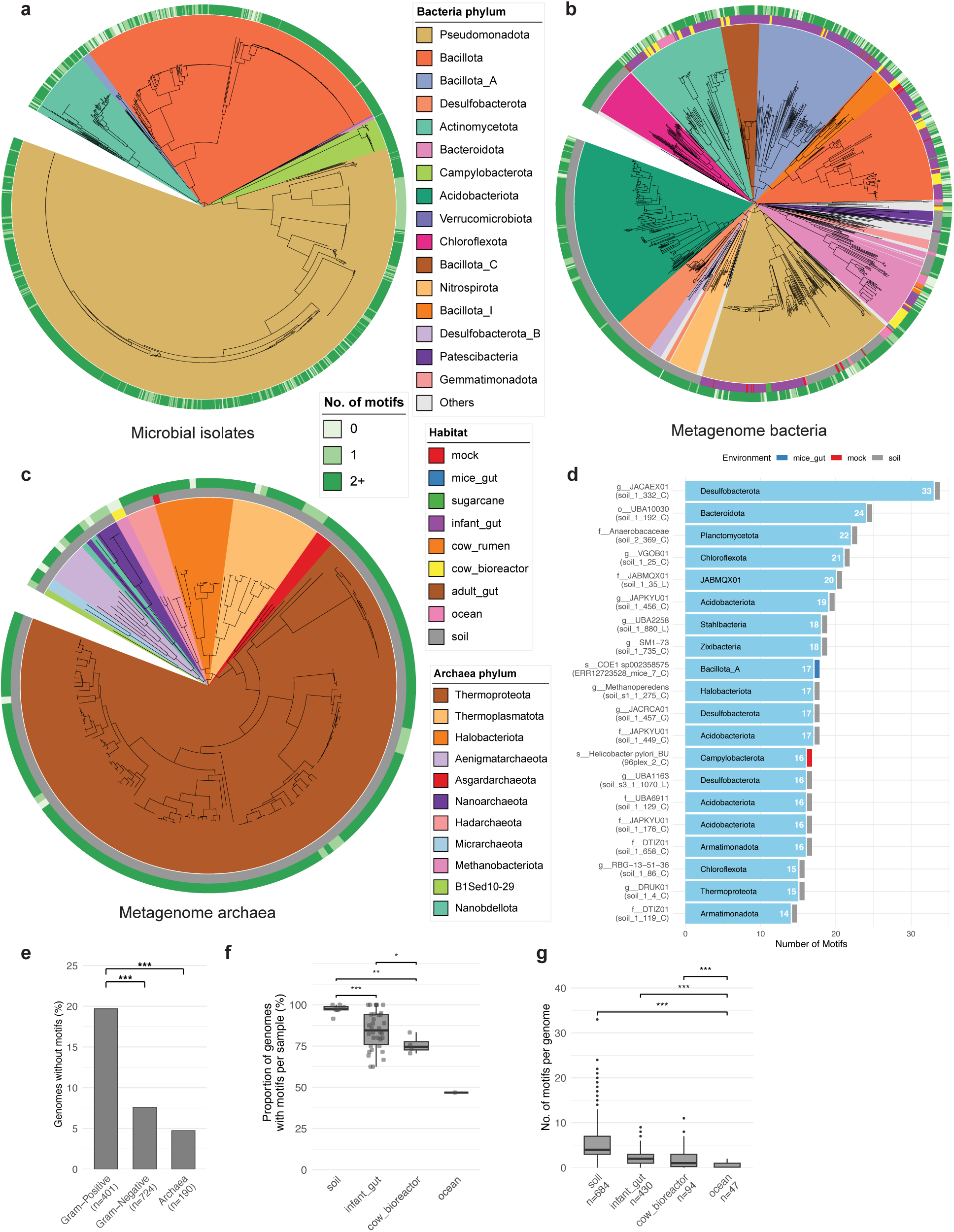
DNA modification landscape across habitats and phyla. **(a–c)** Phylogenetic trees of **(a)** microbial isolates, **(b)** metagenome bacteria, and **(c)** metagenome archaea. Each leaf represents a single genome. Outer ring: Number of motifs detected in the genome. Inner ring: Source habitat of the genome. Branch colors indicate phyla. **(d)** The top 20 MAGs with the highest number of motifs. Left labels indicate the lowest identified taxonomic rank; phyla are labeled inside the bars. The vertical annotation bar on the right indicates the source environment. **(e)** Comparison of the proportion of genomes without motifs between Gram-positive, Gram-negative bacteria, and archaea (Fisher’s exact test). The number of genomes for each category is indicated in the x-axis labels. **(f)** Comparison of proportion of genomes containing motifs per sample, grouped by habitat (t-test). **(g)** Comparison of number of motifs per genome across all samples, grouped by habitat (t-test). The number of genomes for each habitat is indicated in the x-axis labels. For **(e-g)**: Only habitats with no less than 40 genomes are included. Asterisks indicate statistical significance levels: \**P* < 0.05, \*\**P* < 0.01, \*\*\**P* < 0.001.

A strong correlation was observed between the number of motifs and MTases per MAG (Pearson *r* = 0.80, *P*-value = 1.08×10^-291^; Supplementary Figure S6). While genomes typically harbor an average of 3–4 modification motifs (averaging 2.9 in isolates and 4.1 in MAGs)^26,27^, certain genomes exhibit a much higher diversity of motifs. The highest number of motifs detected is 33 in a *JACAEX01* sp. MAG from soil (Figure 2d). Among the 143 unmodified MAGs, 55.24% are Gram-positive bacteria, 38.46% are Gram-negative bacteria, and 6.29% are archaea. By comparing the proportions of modified to unmodified genomes across these lineages, statistical analysis revealed that Gram-positive bacteria exhibit a significantly higher proportion of unmodified genomes compared to both Gram-negative bacteria and archaea (Fisher’s exact test, *P*-values = 4.9×10^-9^ and 4.4×10^-7^, respectively; Figure 2e; Supplementary Figure S7). Consistent with their unmodified status, 93.7% (134/143) of these specific MAGs lacked detectable MTases.

The prevalence of DNA modification varied across habitats. Statistical analysis showed that soil samples had a significantly higher modification proportion, defined as the percentage of motif-containing genomes in each sample, than both infant gut (t-test, *P*-value = 7.19×10^-7^) and cow bioreactor samples (t-test, *P*-value = 2.25×10^-3^) (Figure 2f). In contrast, only 46.81% of genomes from ocean water possessed detectable modification motifs. Additionally, soil genomes contained significantly more motifs (t-test, all comparisons *P*-value ≤ 1.44×10^-19^), whereas ocean genomes harbored significantly fewer motifs than all other habitats (t-test, all comparisons *P*-value ≤ 2.59×10^-9^) (Figure 2g).

### DNA modification exhibits strain-level variation

Analyses of both isolate genomes and MAGs indicate that DNA modification frequently varies at the strain level, as defined by ≥99% average nucleotide identity (ANI) clusters. Among multi-member clusters, motif variation was observed in 64.89% (85/134) of isolate clusters and 24.86% (45/187) of MAG clusters (Figure 3a, c). Crucially, both datasets show a convergence in motif variation frequency as cluster size increases. Variant frequencies plateaued at 95.45% (21/22) for isolate clusters with at least 10 members, and reached 100% for MAG clusters with seven or more members (Figure 3b, d). Figure 3e illustrates this diversity within *Klebsiella pneumoniae* cluster 248_1 from MAGs, which exhibit distinct motif sets among members of the same cluster.

**Figure 3.**
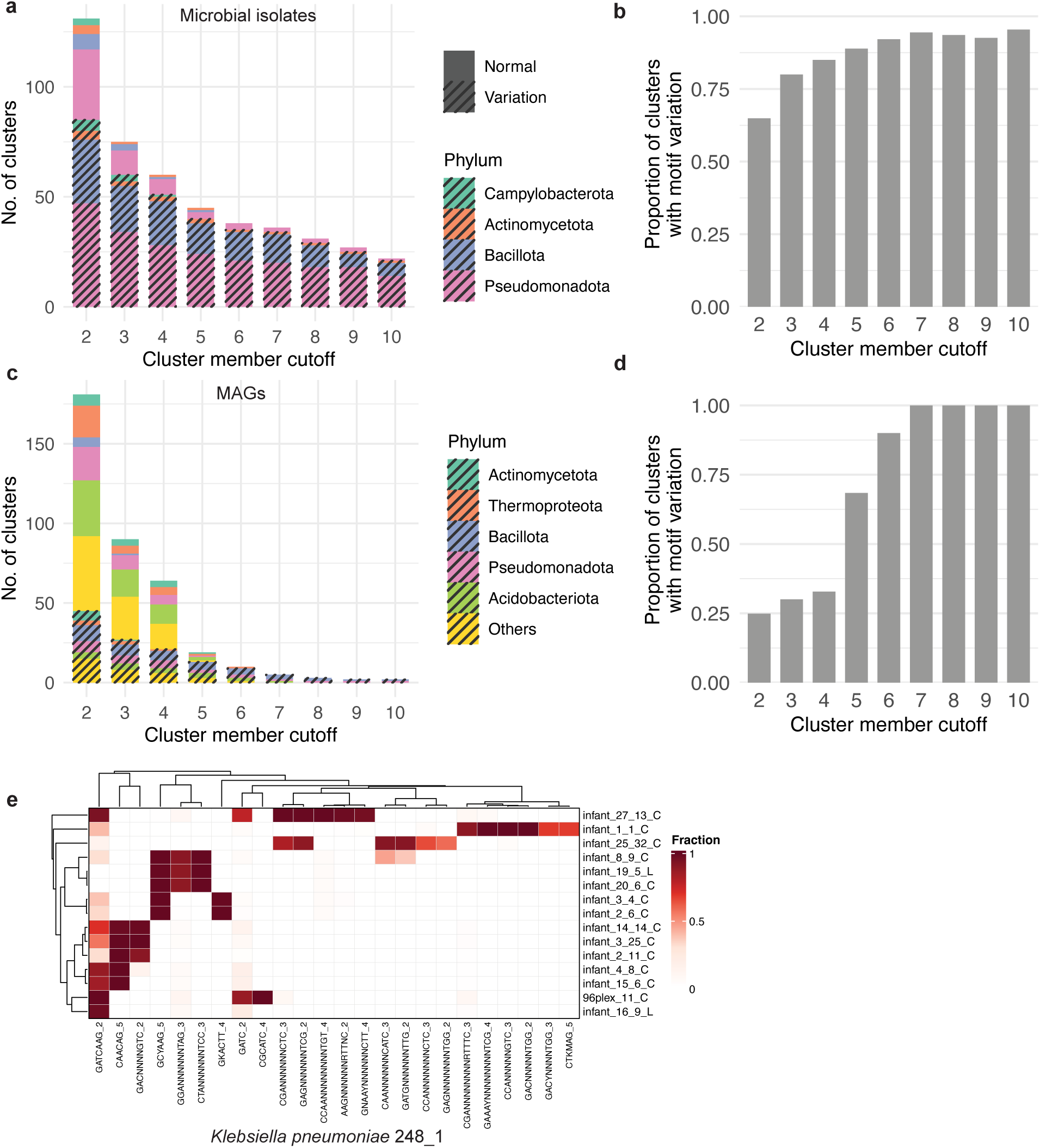
Modification variation within sub-species clusters. **(a, c)** Distribution of motif variation across sub-species clusters (99% ANI). Total number of clusters identified in isolates **(a)** and MAGs **(c)** at varying cluster member cutoffs (2–10). Stacked bars are colored by phylum and subdivided by variation status: solid fills represent "Normal" clusters (consistent modification), while diagonal hatching denotes "Variation" (heterogeneous modification within the cluster). **(b, d)** Proportion of sub-species clusters exhibiting motif variation in isolates **(b)** and MAGs **(d)** across increasing cluster member cutoffs. **(e)** Modification profile of *K. pneumoniae* cluster 248_1 across metagenomic samples. Heatmap of the fraction of modification motifs (x-axis) across longitudinal infant gut samples (y-axis). The color gradient indicates the modification fraction, ranging from 0 (white) to 1 (dark red).

Within each strain cluster, we quantified the relationship between genomic distance and modification similarity by calculating the genome normalized edit distance and modification motif Jaccard similarity for each genome pair (Supplementary Figure S8). We observed that modification similarity is strongly negatively correlated with genome distance in both isolate genomes (Pearson *r* = -0.74) and MAGs (Pearson *r* = -0.73), with *P*-value < 10^-16^ in both cases. Together, these findings demonstrate that DNA modification is highly diverse even within closely related strains.

### DNA modification enables accurate strain-resolved ECE–host linkage inference

We first explored whether ECEs and their hosts share DNA modification motifs by utilizing isolate genomes with established ECE–host linkages. Within these isolates, we identified 1,124 ECEs, comprising 997 plasmids and 127 viruses. The vast majority of these ECEs (89.4%, 1,005/1,124) shared identical motif sets with their corresponding hosts. Although this proportion was relatively low among Bacillota viruses (63.8%, 30/47) (Figure 4a), both plasmids and viruses generally exhibited a high modification Jaccard similarity to their host chromosomes (all ≥ 0.87, Figure 4b).

**Figure 4.**
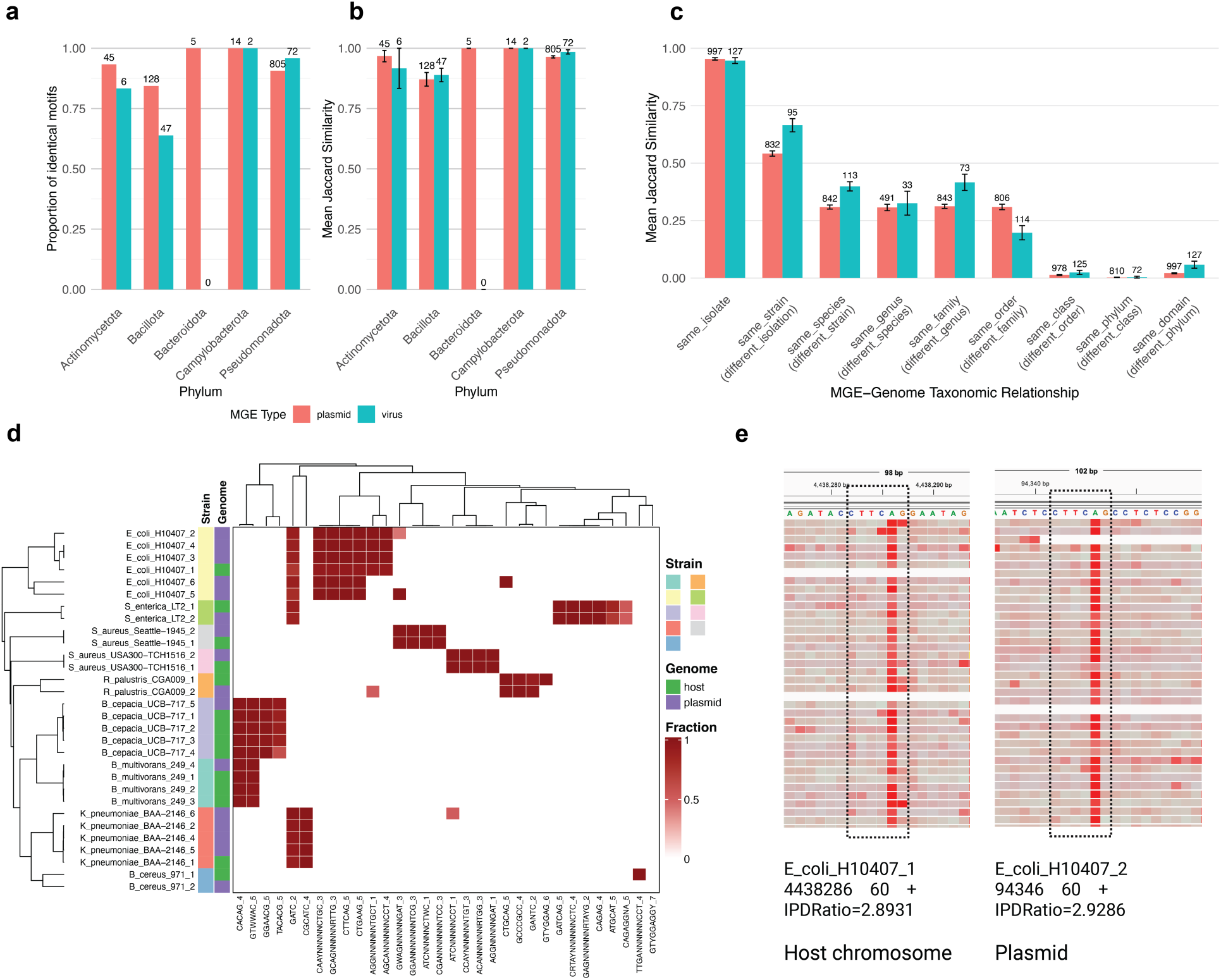
Modification similarity between ECEs and host chromosomes. **(a)** Proportion of ECE–host pairs exhibiting an identical set of modification motifs across various phyla in isolate samples. **(b)** Distribution of ECE–host modification similarity across phyla. Mean Jaccard similarity between the modification profiles of ECEs (plasmids, red; viruses, teal) and their corresponding host genomes in isolates. Numbers above bars indicate the sample size for each category, and error bars represent standard error. Motifs absent from the ECE were excluded in the Jaccard similarity calculation. **(c)** Decay of ECE–genome modification similarity across taxonomic distances. Mean Jaccard similarity of plasmids (red) and viruses (teal) relative to the chromosome across a spectrum of taxonomic relationships. **(d)** Modification profiles of host– plasmid pairs in the PB24 mock community. Heatmap showing modification fractions of motifs across diverse bacterial strains. **(e)** Visualization of IPD values for the motif site CTTC<u>A</u>G in *E. coli* H10407 chromosome and the associated plasmid across reads. Red represents high IPD and white means low IPD, and each block shows a read. Dashed boxes highlighting the consistent IPD peaks at the modification position.

To determine the specificity of this signal, we next examined the impact of taxonomic distance on ECE-genome modification similarity. Modification Jaccard similarity was calculated for ECEs paired with genomes across a spectrum of taxonomic distances, ranging from the exact same isolate to different phyla within the same domain (Figure 4c). Mean similarity peaked for ECE-genome pairs derived from the same isolate (0.95 ± 0.0049) and decreased significantly for pairs originating from the same strain (99% ANI cluster) but different isolates (0.56 ± 0.0107). Similarity further declined for pairs from the same species but different strains (0.32 ± 0.0136), demonstrating that motif sharing is a highly reliable metric for high-resolution linkages. We subsequently validated these biological findings using a PB24 mock community consisting of 24 bacterial strains and 16 known plasmid-host pairs, which demonstrated highly consistent motif sharing between plasmids and their respective hosts (Figure 4d). The modification Jaccard similarity for 15 of these 16 plasmid-host pairs reached 1.0 (Supplementary Figure S9). Illustrating this at the read level, Figure 4e shows a consistent elevation of IPD values at shared motif sites across both the plasmid and host chromosome reads in *E. coli* H10407.

Building upon this foundation, we ran MODIFI on the PB24 mock community to systematically score the confidence of ECE–host linkages. Using subsampled datasets with mean depths ranging from 7× to 140×, linkage recall increased from 43.8% to 81.3% while maintaining 100% precision (Supplementary Figure S10a), with recall reaching a plateau at a mean depth of 44×. Notably, we successfully assigned two plasmids to specific *S. aureus* strains, LT2 and Seattle-1945, and correctly predicted the host strain for five *E. coli* H10407 plasmids even in the presence of the closely related *E. coli* K12-MG1655 strain. Moreover, when evaluating a merged dataset containing both the PB24 data and an infant gut microbiome dataset, MODIFI maintained performance consistent with the mock-only results (Supplementary Figure S10b). Furthermore, using a strain mixture community containing eight species where each species is represented by two distinct strains, we demonstrated that MODIFI reliably links ECEs to their exact host strain (Supplementary Figure S10c and Note S3). In addition, MODIFI exhibited robust linkage estimation across varying community sizes, confirming its scalability (Supplementary Figure S10d–f and Note S3).

Finally, we assessed the tool’s real-world applicability using three cow bioreactor metagenomes where Hi-C was performed in parallel. In these complex datasets, MODIFI-predicted linkages exhibited significantly higher Hi-C contact values than background expectations (mean: 860 vs. 0; t-test, *P*-value = 2.42×10^-2^; Supplementary Figure S10g). Taken together, these stepwise analyses validate the accuracy and high resolution of the ECE–host linkages inferred by MODIFI.

### Large-scale, strain-resolved ECE–host linkage across diverse habitats

Across the 59 metagenomes from nine distinct habitats (Supplementary Figure S3), MODIFI identified 379 high-confidence linkages (linkage score >0.8, specificity <0.001, minimum phylum-level annotation) (Figure 5a; Supplementary Table S5). We observed significant correlations between ECEs and their hosts in both GC content (Pearson *r* = 0.76) and sequencing depth (Pearson *r* = 0.70), as well as a high tetranucleotide cosine similarity (0.95 ± 0.07) (Supplementary Figure S11a-c). Independent validation through CRISPR-Cas spacer matching with zero mismatches further confirmed 4.49% (17) of the linkages (Supplementary Figure S11d). The infant gut metagenomes contained significantly more plasmid linkages than virus linkages, whereas cow bioreactor and soil environments exhibited a non-significant trend toward higher number of virus linkages (Figure 5b). Taxonomically, the phyla Pseudomonadota, Bacillota_A, and Bacillota showed significantly more linkages with plasmids than with viruses (Figure 5c).

**Figure 5.**
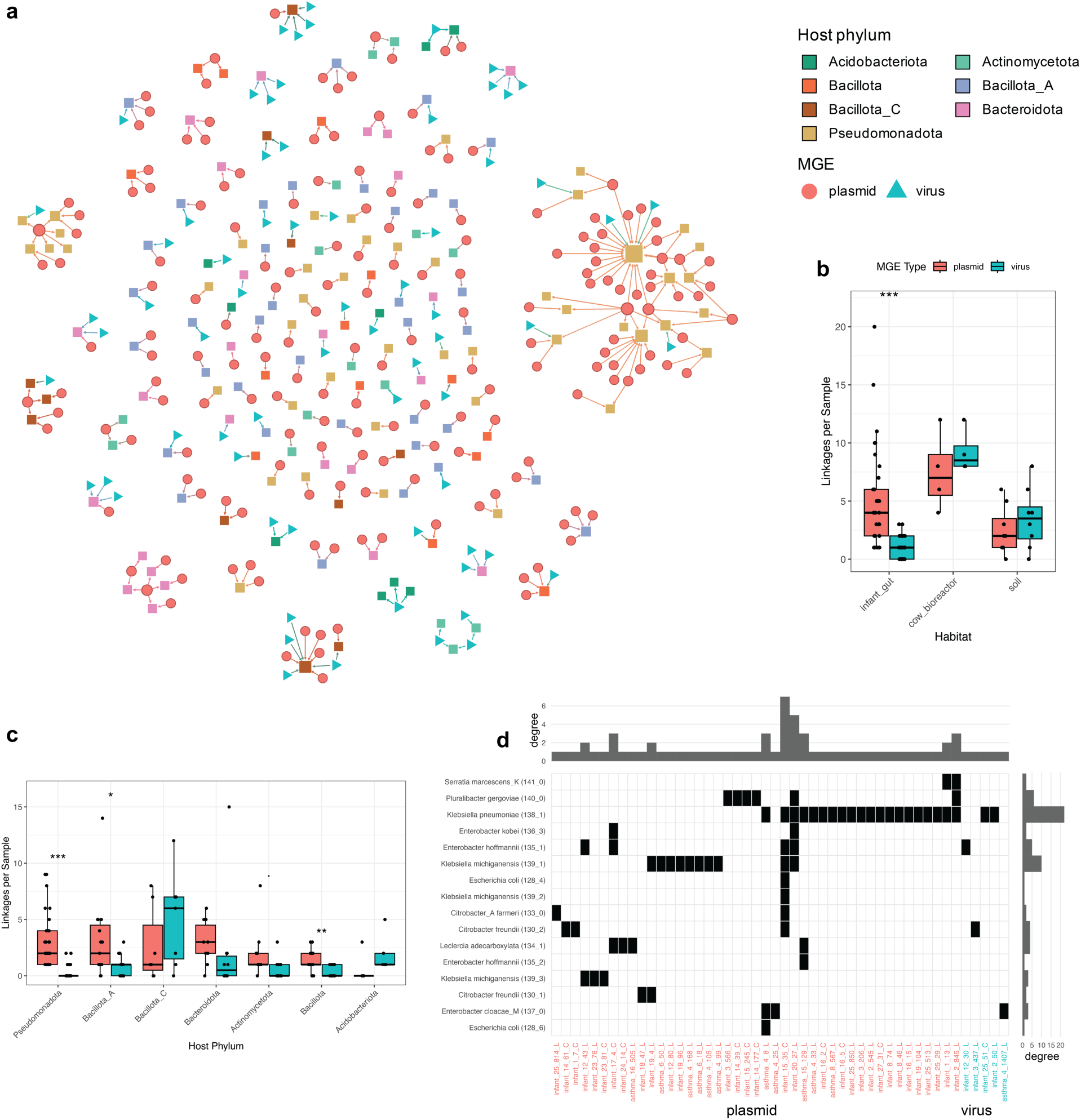
Landscape of ECE–host linkages across diverse habitats. **(a)** Network visualization of inferred host–ECE linkages. Connections are displayed between host genomes (colored squares by phylum) and ECEs, categorized as plasmids (circles) and viruses (triangles). Only the phyla with >5 genomes are shown. Node size represents its degree. **(b)** Abundance of host–ECE linkages by habitat. Boxplots represent the number of predicted linkages per sample for plasmids (red) and viruses (teal) across habitats. Only habitats with at least 4 samples are included. **(c)** Host phylum-specific ECE linkages. The distribution of linkages per sample is shown across top 7 host phyla. Only phyla occurring in at least 5 samples are included. **(d)** Illustration of ECE–host linkages within the largest connected component. The heatmap indicates the presence (black squares) of specific host–ECE linkages, with host subspecies clusters on the y-axis and individual ECEs on the x-axis. ECE types are distinguished by color, where red denotes plasmids and teal denotes viruses. Marginal bar plots summarize the total number of linkages detected per ECE (top) and per host subspecies cluster (right). For **(b-c)**, asterisks indicate statistical significance levels: \**P* < 0.05, \*\**P* < 0.01, \*\*\**P* < 0.001.

To explore these dynamics broadly, we clustered host genomes and ECEs at 99% and 95% ANI, respectively, yielding 315 non-redundant linkages that we rendered as an ECE–host interaction network (Figure 5a). The resulting network consists of 447 nodes, including 174 host, 191 plasmid, and 82 virus clusters. In the network, the largest connected component comprises 16 host, 43 plasmid, and 5 virus clusters (Figure 5d). Notably, all nodes in this major component were recovered from the infant gut, and all host genomes belong to the family *Enterobacteriaceae*. Within this dense subnet, MODIFI successfully achieved strain-resolved ECE linkages for three *Klebsiella michiganensis* clusters and two *E. coli* clusters. *Klebsiella pneumoniae* cluster 138_1 exhibited the highest connectivity, associating with 22 ECE clusters. In this component, plasmid cluster infant_15_35_C linked to as many as seven distinct host clusters, whereas all virus clusters were linked to a single host, consistent with the typically narrow host range of viruses. Taken together, these results highlight the utility of MODIFI for untangling complex, strain-level ECE–host interaction networks across diverse microbial habitats.

### Inversion alters methylation of *Enterococcus faecalis* and associated plasmids in infant gut

Longitudinal analysis of the *Enterococcus faecalis* chromosome and its plasmids revealed a simultaneous methylation shift driven by a site-specific genomic inversion in the infant gut. In infant *I_1*, the methylation motif CA<u>A</u>(N)_6_RTGA predominated from day of life (DOL) 14 to DOL 21 before being replaced by C<u>A</u>Y(N)_6_TAYG from DOL 28 onward (Figure 6a). This transition occurred within a seven-day window and was mirrored in two co-occurring plasmids (59,662 bp and 58,178 bp), which exhibited methylation changes synchronized with the host genome (Figure 6b; Supplementary Figure S12). While this shift was significant for these two bipartite motifs, the ATGC<u>A</u>T motif remained consistent across all four time points spanning DOL 14 to 35.

**Figure 6.**
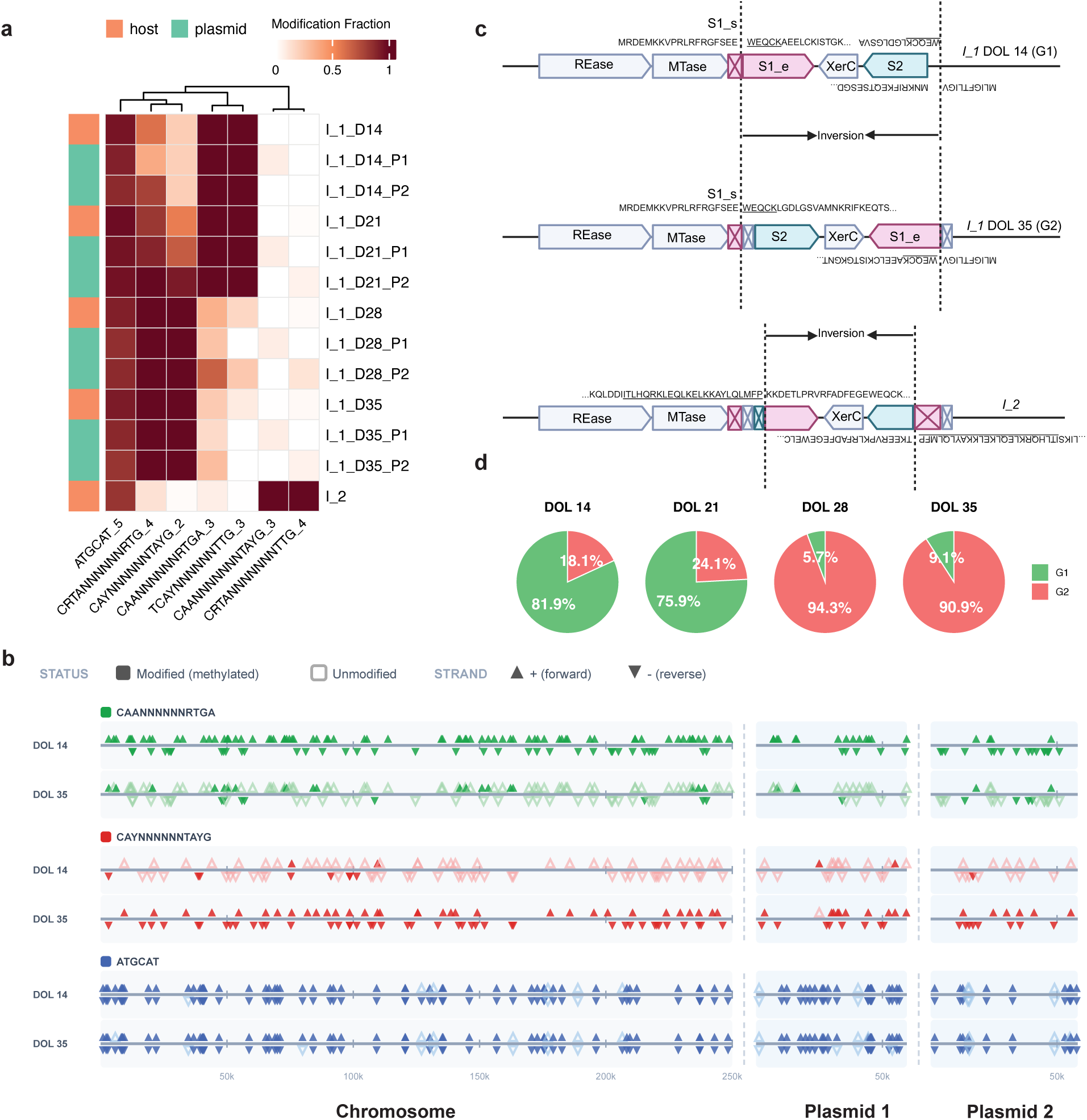
Site-specific genomic inversion drives a synchronized methylome shift in *E. faecalis* and its plasmids. **(a)** Longitudinal tracking of DNA modification profiles across *E. faecalis* populations in infant gut samples, where heatmap colors indicate modification fractions, left sidebars distinguish between host and plasmid genomes, *I_1* and *I_2* represent the first and second infants, *D\** indicates the day of life, and *P\** denotes specific plasmids. Bipartite motifs are shown as reverse-complementary pairs, where CA<u>A</u>(N)_6_RTGA, C<u>A</u>Y(N)_6_TAYG, and CA<u>A</u>(N)_6_TAYG are paired with TC<u>A</u>Y(N)_6_TTG, CRT<u>A</u>(N)_6_RTG, and CRT<u>A</u>(N)_6_TTG, respectively. **(b)** Illustration of shifts in genomic modification sites across a chromosome fragment and two associated plasmids in *E. faecalis* at two time points. At each motif site, filled triangles represent modified loci and empty triangles represent unmodified loci. **(c)** Schematic of the Type I RM system architecture illustrating the inversion across two S subunit genes, with underlined DNA sequences highlighting microhomology at the junction sites. *XerC* is the integrase within the inversion. **(d)** Proportion of two *E. faecalis* genotypes, G1 and G2, across time points of infant *I_1*.

Comparison of complete genomes assembled from DOL 14 and DOL 35 identified a 3.2 kbp inversion as the sole genomic difference (Figure 6c), with the two associated plasmids remaining identical across both time points. This inversion reconfigured two specificity (S) subunits of a Type I RM system designated S1 and S2. At DOL 14, the S1 start codon was located upstream of the inversion point and the gene ended within the inverted region. Following the inversion, the segment of S1 inside the inversion fused with a new sequence to generate a novel N-terminal protein sequence while preserving the original core and C-terminus. Simultaneously, S2 was entirely contained within the inversion at DOL 14, but gained access to a new start codon at DOL 35 to form a hybrid S unit utilizing the original N-terminus of S1 (Figure 6c). The presence of the integrase *XerC* and microhomologies at the inversion junctions suggest a site-specific recombination mechanism. We designated these two genomic configurations as genotypes G1 and G2.

Read alignment analysis confirmed that the DNA modification pattern shift resulted from the rapid expansion of the G2 genotype within the population. Although G2 existed at DOL 14 and 21, it became dominant by DOL 28 and increased from 24.1% to 94.3% of the total *E. faecalis* population within seven days (Figure 6d; Supplementary Figure S13). Similar epigenetic plasticity was observed in a second infant (*I_2*) where *E. faecalis* displayed a distinct CA<u>A</u>(N)_6_TAYG motif (Figure 6a). This variation resulted from an independent inversion event in the S-subunit region, producing different hybrid S-protein sequences (Figure 6c). This result confirms the presence of this site-specific recombination mechanism across independent infant gut environments. Ultimately, this longitudinal analysis demonstrates that MODIFI can uncover dynamic, modification-based population variation, deepening our understanding of rapid microbial adaptation and evolution *in vivo*.

### Borgs and mini-Borgs share modification patterns, but differ from host patterns

Borgs are a novel class of giant archaeal ECEs that remain uncultivated^28–31^. By leveraging MODIFI, we investigated whether Borgs, mini-Borgs, and other ECEs share DNA modification patterns with each other and their hosts of the genus *Methanoperedens*, across eight soil metagenomes (Supplementary Figure S14). Network analysis of modification motif similarities (Figure 7) revealed high modification diversity among *Methanoperedens* species. Viruses, plasmids and mini-chromosome-like elements consistently shared similar modification patterns with *Methanoperedens*. Borgs and mini-Borgs exhibited distinct modification patterns with associated *Methanoperedens*, yet Borgs and mini-Borgs modification patterns were similar. For instance, specific groupings, such as Iris Borg and a Uranus mini-Borg, and Green Borg and two Jupiter mini-Borgs, shared consistent modification patterns within the same samples.

**Figure 7.**
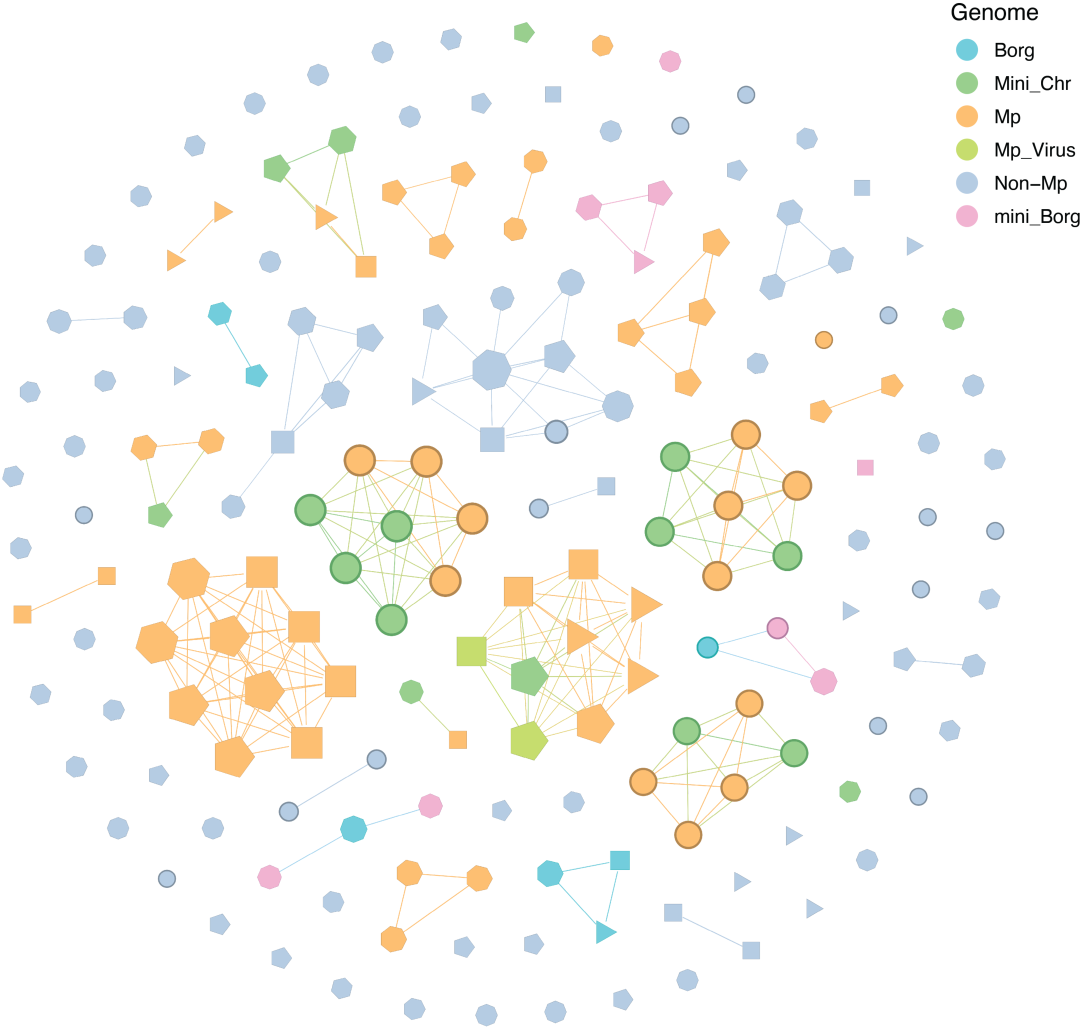
Modification similarity network of *Methanoperedens* and related ECEs. Network visualizing the modification similarity between *Methanoperedens*, related ECEs, and non-*Methanoperedens* genomes. Each node represents a distinct genome, color-coded by its specific type, with edges connecting nodes that share a modification similarity over 0.7. Mp indicates *Methanoperedens* and non-Mp indicates random non-*Methanoperedens* genomes. Mini_Chr represents plasmids and mini-chromosome-like elements. The node shape distinguishes between different soil metagenomes.

We observed striking modification patterns associated with Borg tandem repeats (TRs). Approximately half of all TRs are located within coding regions^32^ (Supplementary Table S6), where they show a significant enrichment for the motif Y<u>C</u>TK^29^ compared to intergenic TRs; this enrichment is significant in 86.36% of diverse Borg types (19/22). In the three Black and one Green Borg genomes with high sequencing depths (≥10×), the modification fraction of Y<u>C</u>TK is significantly higher in intergenic regions with TRs compared to the rest of the genome (Supplementary Table S7). These findings suggest that Borg modification may be involved in gene expression regulation via a TR-related mechanism.

## Discussion

Although the potential for detecting DNA modification using long-read sequences was proposed over a decade ago, current methods remain largely confined to genomes of isolated bacteria^12,16,26,33^. Few studies have attempted to characterize modifications within metagenomes^24,34^, largely due to the lack of scalable tools capable of handling the extreme complexity of metagenomic samples. Ironically, MODIFI leverages the genomic complexity of metagenomes that normally complicates microbiome analysis to enable modification detection without production of unmodified controls. By using an *in situ* self-normalization framework, MODIFI overcomes the primary bottleneck in metagenomic modification detection.

Our findings validate the near-ubiquity of DNA modification across prokaryotic phyla^26^, yet reveal a significantly higher proportion of unmodified genomes in Gram-positive lineages. The robust, multi-layered peptidoglycan cell wall of Gram-positive bacteria provides a formidable physical barrier that can be further reinforced via glycosylation to sterically hinder ECE adsorption. We posit that this superior surface-level exclusion reduces the selective pressure to maintain metabolically costly intracellular immunity, such as RM systems. The envelopes of Gram-negative bacteria and archaea may necessitate a diversified cytoplasmic immune repertoire to degrade invading DNA^35^.

DNA modification motif diversity appears to be governed by habitat-specific ecological constraints, with the marine and soil microbiomes representing opposite ends of the motif diversity spectrum. The motif minimal diversity observed in ocean water may be linked to the evolutionary preference for “genome streamlining” in low-energy-density environments^36–39^. Here, the metabolic cost of maintaining complex RM systems may outweigh the risk of viral predation. Marine microbes may use alternative strategies to reduce viral mortality, such as high strain-level microdiversity and recombination^40^. Conversely, the nutrient-rich, high-density matrix of soil ecosystems supports a diverse repertoire of modifications, likely necessitated by intense horizontal gene transfer (HGT) and viral pressure. Interestingly, while the human gut is typically considered a greater hotspot for defense systems than soil^39^, we observed lower modification diversity in the infant gut compared to soil. This disparity may reflect the low community complexity and immature state of the infant gut microbiome, where narrow niche colonization precedes the establishment of an extensive modification repertoire.

Beyond broad taxonomic trends, we observed strong motif heterogeneity among closely related strains, indicating that DNA modification is a hyper-variable trait at the sub-species level^41^. This strain-level divergence is potentially facilitated by the frequent HGT of defense systems^42^ as has been observed within the *Bacteroides fragilis* group^43^. Furthermore, while phase variation has been previously observed in bacterial genomes^25,44^, our *E. faecalis* strain tracking indicates that this phenomenon occurs simultaneously with the host and its plasmid. The shift indicates that methylation patterns are potential determinants of fitness.

The high concordance between host and ECE modification profiles, coupled with the pervasive strain-level variation of motifs, establishes shared modifications as high-fidelity signatures for mapping *in situ* ECE–host associations. Using network analysis, we inferred *Klebsiella pneumoniae* as a dominant hub of HGT. Given its status as a critical multidrug-resistant pathogen, these linkages provide a framework for tracking the dissemination of antibiotic resistance genes across clinical and environmental populations^45^.

A limitation of MODIFI is that it does not report the specific modification type. However, its scalability allows for the transition from studying isolated laboratory ’type-strains’ to capturing modification variability in complex communities. Although motif matching was effective for ECE– host linkage detection in many cases, it was not universally useful. For example, it is clear that Borgs and mini-Borgs replicate in *Methanoperedens* species^28–31^, yet shared modification patterns were not identified. This may be due to a different mechanism of host interaction and altered functions of modified bases, which were linked to short motifs with relatively low modification incidence. This may point to regulatory functions of modifications, especially when associated with periodic tandem repeats, rather than defense. The method failed to assign one known ECE– host linkage in a mock community and many linkages in metagenomes may have been missed. Lack of linkages may be due to reliance on alternative defense strategies, small ECE genomes (i.e., low incidence of modified bases) and low abundance of ECEs in metagenomes.

MODIFI provides a scalable framework for high-throughput modification discovery across complex microbiomes. The approach will enable analyses of the role of epigenetic modifications in controlling microbial and microbial community activity. MODIFI facilitates large-scale analyses of the strain-level variation in modification patterns, and allows for the recovery of ECE–host linkages directly from PacBio sequencing. By integrating ECE–host linkages into ecological and epidemiological frameworks, MODIFI can enhance our understanding of microbial-ECE interactions^1,2^. Furthermore, the platform identifies potentially compatible vectors and genetic components directly from metagenomic data, providing a critical resource for the precision engineering of microbes^46–49^.

## Methods

### Algorithm of MODIFI

***Read alignment*** PacBio sequencing reads containing polymerase kinetics information (i.e., IPD) are aligned to the metagenome reference using PBMM2 (version 1.14.0) with default parameters^50^. Reads with a mapping quality (MAPQ) score below 20 are discarded. To further ensure alignment reliability, we also apply an identity-based filter. For each read alignment, the alignment identity is calculated as:

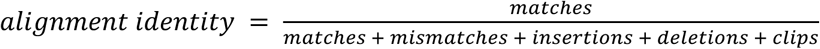

where *mismatches* include only base substitutions, and *clips* refer to the total length of soft- and hard-clipped regions. Only insertions and deletions shorter than 50 bp are considered in alignment identity calculation. While longer insertions and deletions are frequent in metagenomic data due to population-level variation, they do not necessarily indicate poor alignment quality; therefore, they are excluded from the identity filter to avoid penalizing otherwise reliable reads. MODIFI is compatible with both HiFi reads and subreads as input. By default, HiFi reads with alignment identity above 97% and subreads with identity above 85% are retained for downstream analysis. Following alignment, the metagenome was partitioned into individual contig-level FASTA and BAM files to facilitate parallel processing. Contigs shorter than 1,000 bp were excluded from all downstream analyses by default (Figure 1a-c).

### Extraction and processing of IPD signals

To extract polymerase kinetic information, IPD values are loaded from reads alignment per genomic locus. To reduce the influence of anomalous signals, such as those from reads exhibiting globally elevated IPD values, the mean IPD is calculated for each individual read, and the top 5% of reads with the highest per-read average IPD are discarded. Additionally, IPD values equal to zero are excluded from all calculations to avoid artifacts. At each genomic position, the mean IPD value is computed across all remaining aligned reads. For HiFi reads, the Python *pysam* module is used to access aligned kinetic data; for subreads, the Python *pbcore* module is employed. All IPD signals are computed in a strand-specific manner to preserve directional modification patterns. Each genome is divided into 100 kbp chunks for memory management, and all stages of IPD extraction are parallelized using multi-threading.

### Control IPD inference

Since both local DNA context variance and DNA modifications impact IPD values, comparing observed IPD against an unmodified control is essential to detect modifications (Figure 1d). To eliminate the need for sequencing separate control samples that erase all modifications, we propose to estimate the control IPD value directly from the metagenomic data itself. This strategy is based on the observation that most IPD variance is explained by local sequence context (*k*-mer)^21^. Because DNA modifications in prokaryotes are motif-driven and these motifs typically differ between species and strains^14^, our approach assumes that each given *k*-mer will remain unmodified at the majority of its positions within a complex metagenomic reference. This ensures that the mean IPD value of a *k*-mer reflects its unmodified baseline. Supplementary Note S4 and Figure S15 demonstrate that control IPD values derived using this strategy effectively converge to the true unmodified baseline as metagenome length increases.

Following the previous research^21^, MODIFI utilizes *10*-mers by default, with seven bases upstream and two bases downstream of the incorporation site^21^. Specifically, for a base at position 𝑐_0_, the *k*-mer is represented as

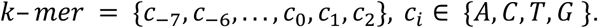

Each *k*-mer is uniquely encoded into a decimal value using a positional hash function:

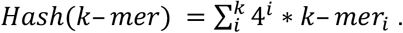

This Hash value serves as an index to store and retrieve *k*-mer-specific information^51^. We traverse the entire assembled metagenome to enumerate all DNA bases and get the *k*-mer of each position. The Hash value of each *k*-mer is used as an index to store the cumulative IPD values and their counts in two separate arrays. After completing the enumeration, MODIFI computes the mean IPD value for each unique *k*-mer.

In a second pass, we re-enumerate each contig. At each base position, we obtain the native IPD value, defined as the average IPD of all reads aligned to that position. We then use the *k*-mer of that position to retrieve its mean IPD value, which serves as the control IPD for that position (Figure 1e). The IPD ratio is calculated as:

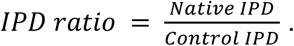

A minimum depth of 3× is required for each locus by default. To optimize performance, this entire step is implemented in C++ using multi-threading to enable efficient processing of large metagenomic datasets.

### DNA modification detection

To detect modified sites, MODIFI assesses positions with significantly elevated IPD ratios (Figure 1e). For each contig, all IPD ratios are collected and modeled as an approximately normal distribution, typically centered around a mean value close to 1. Based on the assumption that most positions are unmodified, we compute the mean and standard deviation of this distribution to define the expected baseline of unmodified positions. Subsequently, for each base position, we calculate a Z-score representing the number of standard deviations its IPD ratio exceeds the mean. From this, we derive a one-tailed *P*-value corresponding to the probability of observing an IPD ratio as higher under the null hypothesis of no modification.

To quantify the confidence of each modification call, we assign a Phred-transformed quality value (QV) to each base, computed as:

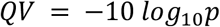

QV values greater than 60 are capped at 60. This score reflects the statistical significance of modification at each base, with higher QV values indicating greater confidence in the detection of a true modification. All detected modified bases are output in GFF format, which includes information for each site such as the QV value, local sequence context (*k*-mer), IPD ratio, and coverage. By default, a QV cutoff of 30 is applied for the following analyses, corresponding to a *P*-value of 0.001.

### Motif detection

To identify *de novo* modification motifs, MODIFI uses MotifMaker from the SMRT Link software suite (version smrtlink-release_25.3.0.273777), with MultiMotifMaker as an alternative^50,52^. Modification motifs are detected by comparing the local sequence context surrounding modified sites across the genome. Subsequently, the motifs are filtered. In each contig, the motifs with modification fraction less than 0.3 and number of modified motif sites less than 100 are removed by default. Modification fraction refers to the number of modified sites divided by the total number of motif strings in a contig.

In regions of low sequencing coverage, modification signals can be noisy, leading to reduced detection sensitivity for modification motifs. In particular, low-depth regions may result in weak enrichment of true motif sequences, thereby compromising motif discovery. To mitigate this issue, MODIFI implements a coverage-based filtering approach. Specifically, we apply a sliding window strategy to identify and mask regions with insufficient read depth (<5×). Only unmasked regions are used for motif discovery to increase the sensitivity of motif discovery.

### Modification profiling for metagenome

Motifs detected across all contigs were aggregated, and the modification fraction for each motif is calculated per contig. To identify all occurrences of each motif within a contig, MODIFI uses the *nt_search* function from the *Bio.SeqUtils* module in *Biopython*. These motif positions are then screened to count the number of modified bases at the identified sites. This yielded the modification fraction of each motif on each contig, which we refer to as the *modification profile* of the metagenome. This approach enables the recovery of modification information for short or low-depth contigs. By leveraging motifs detected in other genomes within the same sample, MODIFI can assign modification status to these motifs even when *de novo* detection is not possible.

### ECE–host linkage inference

To identify hosts for ECEs, we develop a linkage scoring system based on shared DNA modification motifs between ECEs and potential hosts within the metagenome community. We first estimate the motif frequency in the community by calculating length-aware frequencies across the metagenome to account for assembly fragmentation. Additionally, we use a Bayesian Beta-Binomial model to account for bias in low-complexity communities. Based on these frequencies, we quantify the specificity (*S*) of the shared motif set between an ECE and a potential host. A high *S* value indicates that the motif set is common and could occur randomly, whereas a low *S* value suggests a non-random ECE–host association.

Further, we calculate a linkage score (*Φ*) that rewards the specificity of the shared motif set between the ECE and the potential host while penalizing “missing” modification motifs in the ECE that are present in the potential host. This score is weighted by the number of matched motifs and the total number of modified sites. For each ECE, the contig or bin with the highest linkage score is identified as the inferred host. Additionally, we report the GC content and sequencing depth of both the ECE and the inferred host, along with the cosine similarity of their tetranucleotide frequencies. A detailed description of the linkage score calculation framework is in Supplementary Note S5.

### Benchmark and analyses datasets

To validate base modification detection of MODIFI, we used *E. coli* C227 isolate data that included both native and whole-genome–amplified (WGA) control datasets^11^. The performance of modification motif detection was further assessed using the JF8 mock community^14^. We also utilized a PacBio dataset comprising 24 bacterial strains, with each strain prepared as four independent replicates. From these 96 individual whole-genome sequencing libraries, we bioinformatically constructed a mock community designated PB24. This dataset served as a benchmark to evaluate the performance of our modification motif detection and host–ECE linkage inference methods. In addition, three cow rumen bioreactor samples with paired Hi-C data were analyzed to evaluate linkage accuracy. To profile modification in isolates and examine modification similarity between ECEs and their hosts, we retrieved microbial isolates with PacBio HiFi data from NCBI (accession numbers listed in Supplementary Table S2).

For large-scale environmental analysis, we collected 59 PacBio HiFi metagenomic datasets spanning nine habitats, including soil, cow rumen, cow rumen bioreactor, mice gut^53^, infant gut, adult gut, ocean water, sugarcane^54^ and PB24 mock community samples (Supplementary Table S3). The mouse gut and sugarcane sequences were sourced from previously published datasets. The soil, cow rumen, cow bioreactor, and infant/adult gut ^55^ datasets were generated by our group and collaborators. To ensure all analyses were based on HiFi reads, the study only included data generated by PacBio Sequel II/IIe and Revio platforms. PacBio subreads were processed into HiFi reads using the SMRT Link ccs tool^50^ with the following parameters: *--hifi-kinetics --min-rq 0.99 --min-passes 3*.

### DNA extraction, sequencing, and processing

For the *in vivo* cow rumen and *in vitro* rumen bioreactor datasets, 3 samples were subject to in-house DNA extraction by using DNeasy PowerMax Soil Kit (Qiagen) and another 2 samples were shipped to the genome center in Maryland University to perform high molecular weight DNA extraction. All the DNA samples were sequenced on the PacBio Revio system. For Hi-C data, proximity ligation was performed using ProxiMeta Hi-C kits (Phase Genomics), followed by sequencing on the Illumina NovaSeq platform.

Human gut samples were obtained from two distinct infant cohorts: NICU Antibiotics and Outcomes (NANO) Trial (Clinical Trial ID: NCT03997266)^55,56^ and the 12-month timepoint from the Trial for Infant Probiotic Supplementation (TIPS)^57^. For the NANO cohort, DNA was extracted using the ZymoBIOMICS DNA Miniprep Kit, while for the TIPS cohort, a CTAB-chloroform extraction was used as previously published^58^. Both cohorts were sequenced on the PacBio Revio system.

Soil datasets were obtained from two cohorts. For the first cohort with two samples, DNA was extracted using the DNeasy PowerMax Soil Kit (Qiagen) and sequenced on the PacBio Sequel II system. For the second cohort, six soil samples were collected from varying depths (60–115 cm). Genomic DNA was extracted using a modified DNeasy PowerSoil protocol (Qiagen) designed to minimize shearing for high-molecular-weight DNA. SMRTbell libraries (Prep Kit 3.0) were prepared from 3 μg of sheared DNA per sample, size-selected (>3 kb), and multiplexed for sequencing on the PacBio Revio system, yielding >2.5 million HiFi reads per sample.

### Genome assembly

PacBio HiFi reads from both metagenomic and isolate samples were processed through automated Snakemake workflows. Raw BAM files were converted to FASTQ using *bam2fastq* (SMRT Link^50^) after indexing with *pbindex*, and low-quality bases were trimmed with BBDuk v39.01 (*qtrim=rl, trimq=20, minavgquality=20*). For metagenomic samples, host reads were removed by mapping to the host reference genomes (Supplementary Table S4) with minimap2 v2.28-r1209 (*-ax map-hifi*)^59^, and the host-depleted reads were assembled using hifiasm_meta v0.3-r074 (*--no-binning*)^60^. For isolate genomes, assemblies were generated with hifiasm v0.25.0-r726 (*--primary -l0*)^61^. In both cases, contigs were extracted and renamed from the GFA output using a custom Python script. To minimize ambiguity, all subsequent analyses focused on assembled contigs independent of metagenomic binning.

Genome completeness and contamination were assessed with CheckM2 v1.0.1^62^, and taxonomic classification was performed using GTDB-Tk v2.4.0^63^. Only isolates with contamination levels of 5% or less are considered pure and retained for downstream analysis. Phylogenetic trees were constructed using GTDB-Tk under default parameters and visualized using iTOL^64^. The circos figures illustrating the modification profiles were generated using Proksee^65^. ECEs were predicted with geNomad^66^ v1.7 (*--relaxed --enable-score-calibration --sensitivity 7.0*). Pro-viruses were ignored in all the analyses.

From the assembled metagenomes, we retained high-quality contigs that met at least one of the following criteria: ≥50% completeness with ≤5% contamination, or circularity. Additional filtering required contigs to be not annotated as ECEs, to have at least phylum-level GTDB taxonomic assignment, and to exhibit ≥10× sequencing coverage.

### Genomic strain clustering

To examine whether DNA modification motifs vary within the same strain cluster, we performed independent clustering for the metagenomic contigs and the isolate genomes. In both cases, clustering was conducted using dRep v.3.5.0 with Skani at thresholds of 99% ANI and 70% alignment fraction^67^. Each resulting cluster was treated as a strain.

### Motif counting and modification diversity quantification

To count the number of motifs in each genome, we filtered the motifs with the criteria: min_frac = 0.3, min_sites = 100. This conservative site threshold was selected to minimize false positives and noise. Reverse-complementary motifs were treated as a single entity and counted only once. Moreover, mean motif numbers for isolates and MAGs were estimated using only near-complete genomes (>90% completeness) to minimize potential bias from missing genomic regions.

In strain-level modification diversity analysis, to prevent redundant comparisons of contigs from the same isolate, we selected the longest contig (>500 kb, ≥10× coverage, and not annotated as an ECE) from each isolate to serve as a representative isolate contig. For each multi-genome cluster, to assess the variation in DNA modification, we defined the cluster motif set as motifs with a modification fraction >50% and ≥500 detected loci in at least one genome. Within a cluster, a genome was considered to lack a motif if its modification fraction was <10%.

### Quantify modification similarity and genomic divergence

To quantify the modification similarity between two genomes, we employed two distinct methods: the identification of identical motif sets and the calculation of the Jaccard similarity. In examining whether ECE and host share DNA modification motifs, the ECEs with length < 5 kbp or depth lower than 10x were discarded. To calculate Jaccard similarity between ECE–host pairs, we filtered out host motifs that lacked corresponding target sequences within the ECE. Jaccard similarity was computed by binarizing motif frequencies based on a modification fraction threshold of 0.3.

To quantify genomic divergence between two genomes within the same strain, we performed a comparative analysis using MUMmer3 *dnadiff*^68^. The total edit distance was subsequently calculated as the sum of the total number of SNPs and the total number of InDels. Furthermore, the edit distance was normalized by the length of the shorter genome in each pairwise comparison to account for variation in sequence scale.

### ECE–host linkage validation

To benchmark MODIFI for ECE–host linkage, we analyzed the PB24 mock community consisting of 24 bacterial strains and 16 known plasmid–host linkages. Only MODIFI-inferred linkages with a linkage score exceeding 0.5 and a specificity value below 0.01 were retained for further analysis. Through a subsampling analysis (5%, 10%, 20%, 30%, 50%, and 100% sampling rates), we calculated the mean contig depths for each subsample. To assess performance within more complex metagenomic contexts, we *in silico* amended the subsampled PB24 data into an infant gut microbiome dataset. Further, we merged isolate data to simulate strain-mixture communities to assess the strain-level ECE–host linkage performance of MODIFI (Supplementary Note S3). We also generated mock communities with varying numbers of species to assess how community size impacts linkage accuracy (Supplementary Note S3).

Three cow rumen bioreactor samples with both HiFi and Hi-C data were analyzed to evaluate linkage accuracy. Because Hi-C contact frequencies reflect the physical proximity of DNA sequences within a cell, they provide an independent measure of ECE–host linkage. Hi-C contact matrices were generated using Bin3C v0.1.1a^69^, and contact frequencies were extracted for each ECE–host pair inferred by MODIFI. ECEs with only self-contacts or low total Hi-C contact counts (< 20) were excluded from analysis. For each sample, the Hi-C contact values supporting MODIFI-inferred ECE–host pairs were compared against a background of 1,000 randomly selected contig pairs to assess linkage accuracy.

### CRISPR-Cas spacer matching

CRISPR spacers were predicted from assembled genomes using MinCED 0.4.2 with default parameters^70^. ECEs were identified and extracted from the same assemblies for use as potential protospacer targets. To identify CRISPR-mediated targeting of ECEs, spacer sequences were aligned against the ECE database using BLASTN 2.16.0+ (task: blastn-short)^71^. Raw BLAST hits were filtered using a custom pipeline requiring no mismatches.

### RM gene annotation

RM systems encoded in the assembled contigs were annotated using the MicrobeMod v1.1.0 toolkit^72^, which is specifically designed for the identification of REases and MTases in prokaryotic genomes. The *annotate_rm* module was run with default parameters, using the assembled contig FASTA files as input. The tool searches for homologs of known RM system components by comparing input sequences against curated reference databases of MTase and REase protein families. In the correlation analysis of MTase and motif counts, we quantified MTases by treating each operon as a single functional unit. If an operon contained two MTases, they were counted together as one functional representative for that locus.

### ECE–host linkage network analyses

MODIFI was used to call ECE–host linkages from all the 59 metagenomes from nine habitats. Based on the mock community benchmarks, we initially observed that a linkage score > 0.5 and specificity < 0.01 provided reliable detection. However, to further minimize potential false positives in complex, real-world metagenomic datasets, we applied more stringent empirical thresholds (linkage score > 0.8 and specificity < 0.001) for the final analysis. Additionally, we required that all host genomes possess at least a minimum phylum-level annotation according to the GTDB.

Also, to remove redundancy in the ECE–host linkages detected by MODIFI, we clustered host and ECE genomes. ECE sequences from all samples were first merged into a single FASTA. We built a BLAST nucleotide database from this set and ran an all-vs-all MegaBLAST search of the merged sequences against the same database^71^. Pairwise ANI was computed from the BLAST results, and sequences were clustered using a minimum ANI threshold of 95% and a minimum fractional alignment length of 85%. Host genomes were clustered separately using a 99% ANI threshold.

To construct the linkage network, nodes were defined as either ECE or host clusters, with edges representing the identified linkages between them. Network analysis, including the identification of connected components and the calculation of network metrics, was performed using the Python *NetworkX* module.

### Borg and mini-Borgs analyses

To systematically analyze modifications in *Methanoperedens* and related ECEs, we selected corresponding contigs from the soil metagenomes and included randomly selected non-Methanoperedens genomes as a background. For this genome set, we identified all motifs and calculated the modification fraction for each motif across every genome. Modification similarity was then determined using the Pearson correlation coefficient of these modification fractions. To construct a modification similarity network, genomes were represented as nodes, with edges connecting pairs that exhibited a high modification similarity (Pearson *r* > 0.7). The resulting network was visualized using Gephi^73^. Furthermore, TRs in Borg genomes were identified using a previously reported script^32^, and genome annotation was performed using PRODIGAL v2.6.3^74^ with default parameters. To assess the enrichment of motif sequences or higher modification fractions, we employed Fisher’s exact test.

## Data availability

Microbial isolates were retrieved from NCBI (accession numbers listed in Supplementary Table S2). Data sources of public datasets for the PB24 mock community, JF8 mock community, *E.coli* C227, ocean water, mouse gut, and sugarcane were listed in Supplementary Table S3. The soil, cow rumen, cow bioreactor, and infant/adult gut datasets were generated by our group and collaborators. The raw data supporting the findings of this study will be made available in a public repository upon publication.

## Code availability

The source code of MODIFI is freely available at https://github.com/sachdevalab/MODIFI. The scripts for reproducing the results are also available in this repository.

## Acknowledgements

We gratefully acknowledge the researchers who contributed public isolate and metagenomic PacBio HiFi data. We thank Pacific Biosciences for providing the sequencing services for six soil samples analyzed in this study, as well as for publishing the PB24 mock community dataset. We also acknowledge PacBio and SeqCenter for the 2024 PacBio Microbial Genomics SMRT grant (to A.K.G.) that funded the sequencing of the NANO infant samples. We thank BioRender.com for the assistance in creating the schematic illustrations. We thank Dr. Jaymin Patel, Dr. Siqi Yang, and Colin Robinson for insightful discussions. We thank Leylen Miloslavich, Dr. Shufei Lei, Tasha Kayatsky, and Glen Otero for laboratory and technical support.

## Author Contributions

R.S. and J.F.B. conceived the study. S.W., R.S., and J.F.B. designed the study. A.K.G., M.J.M., and J.F.B. contributed to the NANO infant dataset. L.V.A., M.C.S., R.S., and J.F.B. contributed to the soil datasets. R.E.G., P.Z., M.H., and S.D. contributed to the Hi-C and PacBio data for the cow rumen and bioreactors. H.M.S., P.Z., S.L., and S.D. contributed to the TIPS dataset. D.M.P., S.Z., and J.E.W. performed the PacBio HiFi sequencing for the soil metagenome subset. S.W. and R.S. collected the public data. S.W. and R.S. led software development and modification analyses. S.W., L.-D.S., R.S., and J.F.B. performed the Borg analyses. M.Y. assisted with genome annotation. S.W., R.S., and J.F.B. wrote the manuscript with input and revisions from all authors.

## Supplementary Notes

### Note S1 | Accurate DNA modification detection

To evaluate MODIFI for DNA base modification detection, we analyzed the *E. coli* C227 isolate with both native genomic DNA and WGA control DNA data^11^. Because the data were generated on the PacBio RS II platform, they were incompatible with modern SMRT Link tools for HiFi conversion; consequently, subreads were processed directly for downstream analyses. We constructed a *k*-mer-based empirical control IPD database from the WGA data to analyze the native dataset. The CTGC<u>A</u>G motif was assumed to be fully methylated. Recall was defined as the fraction of methylated CTGC<u>A</u>G sites detected in native DNA, while the false positive rate (FPR) represented detections in control DNA. At 10x coverage, MODIFI achieved 99.47% recall and 0.14% FPR using a *P*-value cutoff of 0.001 (QV=30). Increasing the coverage threshold to 20x further improved performance to 99.58% recall and 0% FPR, demonstrating high sensitivity and specificity of MODIFI (Supplementary Figure S1a).

To assess DNA modification detection in a metagenomic context and validate the *k*-mer-based control IPD estimation method, we simulated two metagenomes by spiking soil metagenomic reads (1% subsampled, 6.2 Gbp DNA bases total) with either WGA or native E. coli C227 reads. We applied MODIFI using default settings, employing the tool’s internal control estimation on each dataset. The estimated control IPD values exhibited a high correlation with the ground-truth values (Spearmon *r* = 0.70, *P*-value < 10^-16^). Across various coverage thresholds, MODIFI maintained robust performance; at 10x coverage, recall was 99.42% with a 0.21% FPR at a *P*-value cutoff of 0.001 (QV=30), while at 20x coverage, recall reached 99.54% with a 0.10% FPR (Supplementary Figure S1b). These results demonstrate that MODIFI accurately detects base-level DNA modifications in complex metagenomic samples with high sensitivity and low error rates.

Furthermore, to validate the motif detection performance of MODIFI, we applied it to the JF8 mock community, which consists of eight distinct species previously reported to possess unique methylation motif profiles^14^. As the data were also generated on the PacBio RS II platform, we performed MODIFI in subreads mode. The motifs recovered by MODIFI were manually compared with previously reported motifs. We recovered 93.75% (15/16) of the previously reported motifs (Supplementary Table S1). The resulting modification motifs clearly distinguish contigs from the eight species (Supplementary Figure S1c).

In addition, to assess the impact of sequencing depth on motif detection of MODIFI, we subsampled the PB24 mock community across varying depth levels. MODIFI was performed on these datasets in HiFi mode with default settings. For each subsample, we quantified the motifs identified and their corresponding relative depths. To normalize these results, we calculated the detection recovery rate, which is defined as the percentage of motifs detected at a given depth relative to the total motifs identified at >50x coverage. As shown in Supplementary Figure S1d, the mean detection percentage scales with depth: at a mean depth of 7x, 66% of motifs are recovered, rising to 94% at 14x. These results indicated that while low depth can lead to missing motifs, MODIFI remains highly effective in depth-constrained scenarios, recovering two-thirds of the motifs even at 7x depth. Overall, these results suggest that MODIFI is reliable for both DNA base modification detection and motif discovery.

### Note S2 | Computational resource consumption evaluation

We first benchmarked the computational efficiency of MODIFI against ipdSummary. Since ipdSummary is incompatible with HiFi reads, we utilized a soil metagenome subreads sample for this comparison. Furthermore, because ipdSummary failed to process the entire metagenome as a single reference, we randomly selected genomic subsets of approximately 100, 200, 300, and 500 Mb for the benchmark. Given the reads alignment on these genomic subsets, both MODIFI and ipdSummary (smrtlink-release_25.3.0.273777) were executed using default parameters on a high-performance computing cluster with dedicated node-level resource allocation to ensure an equitable comparison. As ipdSummary is not designed for motif discovery, the comparison was limited to base modification detection performance.

The results demonstrate that MODIFI consumes significantly fewer computational resources than ipdSummary (Supplementary Figure S2). For a 200 Mb reference, MODIFI completed the analysis in only 2.7 hours of wall-clock time compared to 39.7 hours for ipdSummary—a nearly 15-fold increase in processing speed. This efficiency is further underscored by CPU time requirements: MODIFI processed a 500 Mb reference in 194.8 hours, whereas ipdSummary required more than triple that time (712.3 hours) for a reference only one-fifth that size (100 Mb). Notably, ipdSummary failed to complete the 300 Mb and 500 Mb benchmarks. Furthermore, MODIFI’s memory management maintains a stable peak footprint of 15–18 GB in these tasks. In contrast, ipdSummary exhibited a sharp, non-linear surge in memory consumption, reaching 205 GB for a 200 Mb reference. These trends indicate that MODIFI is suited for large-scale metagenomic applications where ipdSummary and ipdSummary-based tools become computationally prohibitive due to escalating hardware demands and impractical execution times.

In addition, we evaluated the computational resource consumption of MODIFI for integrated base modification detection and motif discovery across 59 PacBio HiFi metagenomic samples from nine habitats (Supplementary Figure S3-S4). Modification detection was performed using default parameters with 64 threads, taking aligned BAM files as input. The mean wall-clock time was 4.5 hours (range: 0.2–19 hours, std: 3.9), while the mean CPU time was 120 hours (range: 1–1,161 hours, std: 215), with a mean peak memory usage of 18 GB (range: 0.6–62 GB, std: 14). The maximum CPU time and wall-clock time were 1,161 hours and 19 hours, respectively, both recorded for the ocean sample, while the maximum peak memory of 62 GB was observed for a soil sample. These results demonstrate that MODIFI is a computationally efficient method capable of handling complex environmental datasets.

### Note S3 | Evaluation of ECE–host linkage inference using mixed isolate communities

To evaluate MODIFI’s ability to resolve strain-level ECE–host linkages, we constructed a mock community, designated StrainMix, by pooling isolate data. We selected eight species, each represented by two distinct strains belonging to separate 99% ANI clusters. Assembled genomes and HiFi reads of these isolate data were merged independently to simulate a metagenomic dataset prior to analysis with the MODIFI pipeline. In total, 27 ECE–host linkages (comprising 22 plasmids and 5 viruses) served as the ground truth for assessing linkage accuracy of MODIFI. At a linkage score cutoff of 0.3, MODIFI achieved 100% recall with 92.6% precision (Supplementary Figure S10c). The two false positives resulted from a failure to distinguish between the two specific strains of *Streptococcus pneumoniae*. Increasing the threshold to 0.38 improved precision to 95.8%, with recall remaining high at 92%. Under a more stringent cutoff of 0.68, MODIFI achieved 100% precision, though recall decreased to 44%. These results demonstrate that MODIFI can effectively resolve ECE–host linkages even between closely related strains within the same species.

To benchmark the scalability of MODIFI’s linkage performance, we constructed five mock metagenomic communities containing 10, 20, 30, 40, and 50 bacterial isolates, each representing a distinct species. These communities carry between 11 and 75 ECEs in total (Supplementary Figure S10d). MODIFI identified these linkages with consistently high recall across all community complexities. At a linkage score of 0, recall exceeded 90% in four of the five communities, with the 10-species community reaching 100% recall prior to any false positive detection (Supplementary Figure S10e, f). While recall declined gradually as the score threshold increased, this trend remained consistent across all community sizes, indicating that MODIFI’s performance does not significantly degrade with increasing community complexity. Precision reached 100% for all communities at a score threshold of approximately 0.45–0.50. Collectively, these results demonstrate that MODIFI provides reliable ECE–host linkages across communities of varying taxonomic breadth and composition.

### Note S4 | Analysis of the control IPD inference

To assess the reliability of MODIFI’s control IPD inference, we performed a theoretical simulation. Consider a modified microbial genome of length *t* within a metagenome of total genome length *X*. For a *k*-mer of length *k*, as 𝑘 ≪ 𝑡 and 𝑘 ≪ 𝑋, the number of *k*-mer occurrences in the modified genome is approximately *t*, while the total occurrences in the metagenome is approximately *X*. There are 4^k^ potential distinct *k*-mers. Assuming *k*-mers are uniformly distributed across genomes, the expected occurrence of each *k*-mer is 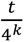 in the modified genome and 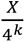 in the metagenome. Suppose a specific *k*-mer is 100% modified in a genome of length *t* and this modification is unique to that genome. Consequently, the average IPD in the metagenome, denoted as *a*, for that *k*-mer can be expressed as

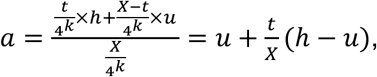

where *h* and *u* represent the modified and unmodified IPD values, respectively. In the MODIFI framework, *a* is regarded as the estimated unmodified IPD value of that *k*-mer. To quantify the deviation of this estimate from the ideal unmodified mean *u*, we define the control IPD deviation σ as:

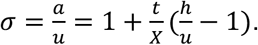

We denote 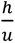 as the modification effect ratio, which represents the factor by which the modified IPD deviates from the control IPD. A σ value closer to 1 indicates a more reliable estimation of the control IPD value.

Given a representative microbial genome length of 5 Mbp, total metagenome lengths ranging from 5 Mbp to 3,000 Mbp, and modification effect ratios of 1.5, 2, 2.5, and 3, the resulting control IPD deviations decrease sharply with increasing metagenome length and eventually converge to 1 (Supplementary Figure S15). At *X* = 100 Mbp, the control IPD deviations range from 1.025 to 1.10. Empirically, these results suggest that a metagenome length of 100 Mbp enables reliable control IPD estimation by minimizing the influence of modified sites on the global *k*-mer mean. These simulation results demonstrate that control IPD inference becomes increasingly reliable as metagenomic complexity increases, further establishing MODIFI as a robust framework for metagenomics.

### Note S5 | Detailed mathematical framework for ECE–host linkage

To predict hosts for ECEs, MODIFI calculates a linkage score based on shared DNA modification motifs. The linkage score quantifies the uniqueness of shared modification motifs between each ECE and potential host in the metagenome (Figure 1f). To quantify the uniqueness of each motif, we calculate its frequency within the metagenome. Since single genomes are often fragmented into multiple contigs during assembly, we define a length-aware motif frequency, calculated as the cumulative length of contigs containing the motif divided by the total length of all contigs in the metagenome. To account for instances where low metagenomic complexity might lead to biased frequency estimates, we utilize a Beta-Binomial model. Let the motif frequency follow a prior distribution Beta(*α, β*) with *α* = 2×10^6^ and *β* = 1.8×10^8^. Given the total length of contigs containing the motif (𝐿_𝑚_) and the total length of all contigs (𝐿_𝑎_), the posterior distribution of motif frequency is:

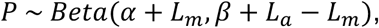

and the posterior mean is:

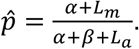

We then use the posterior mean as the point estimate for the motif frequency. For a set of *n* motifs, we estimate the probability of observing the set by chance in the metagenome as:

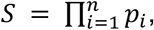

where 𝑝_𝑖_ is the frequency of motif *i*, and *S* represents the specificity of the motif set. A high *S* value indicates that the motif set is common and could occur randomly, whereas a low *S* value suggests a non-random association and therefore stronger host–ECE specificity.

As an ECE may associate with multiple hosts, it can carry the combined set of motifs from all of them, potentially exhibiting more motifs than any single host. To determine whether an ECE and a specific host are linked, we evaluate only the host’s motif set. We scan the ECE genome to calculate modification fractions of host motifs, discarding motifs with fewer than 5 occurrences in ECE. A motif is considered present in the ECE if its modification fraction exceeds 0.3. For each ECE, specificity is calculated based on the shared motifs with each non-ECE contig (or bin). We then define a linkage score *Φ* to identify the most likely host. For motif *i*, define an indicator 𝛾_𝑖_ ∈ {0, 1} for presence in ECE. Let *n* be the number of motifs. The probability of concordance between the host and ECE motifs is:

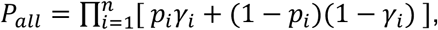

where 𝑝_𝑖_ is the frequency of motif *i*. Normalized by the number of motifs:

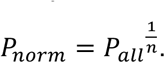

We also track the number of matched motifs (*t*) and total modified sites (*h*). The raw linkage score is then:

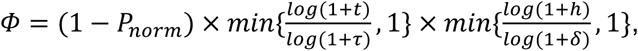

with default values *τ* = 3 and *δ* = 5,000. Finally, we penalize motifs expected in the host but absent in the ECE. Let 𝑓_{ℎ,𝑖}_and 𝑓_{𝑚,𝑖}_be the modification fractions in the host and ECE in the motif *i*, respectively. The penalized linkage score is:

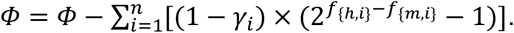

## SUPPLEMENTARY FIGURES

**Figure S1.**
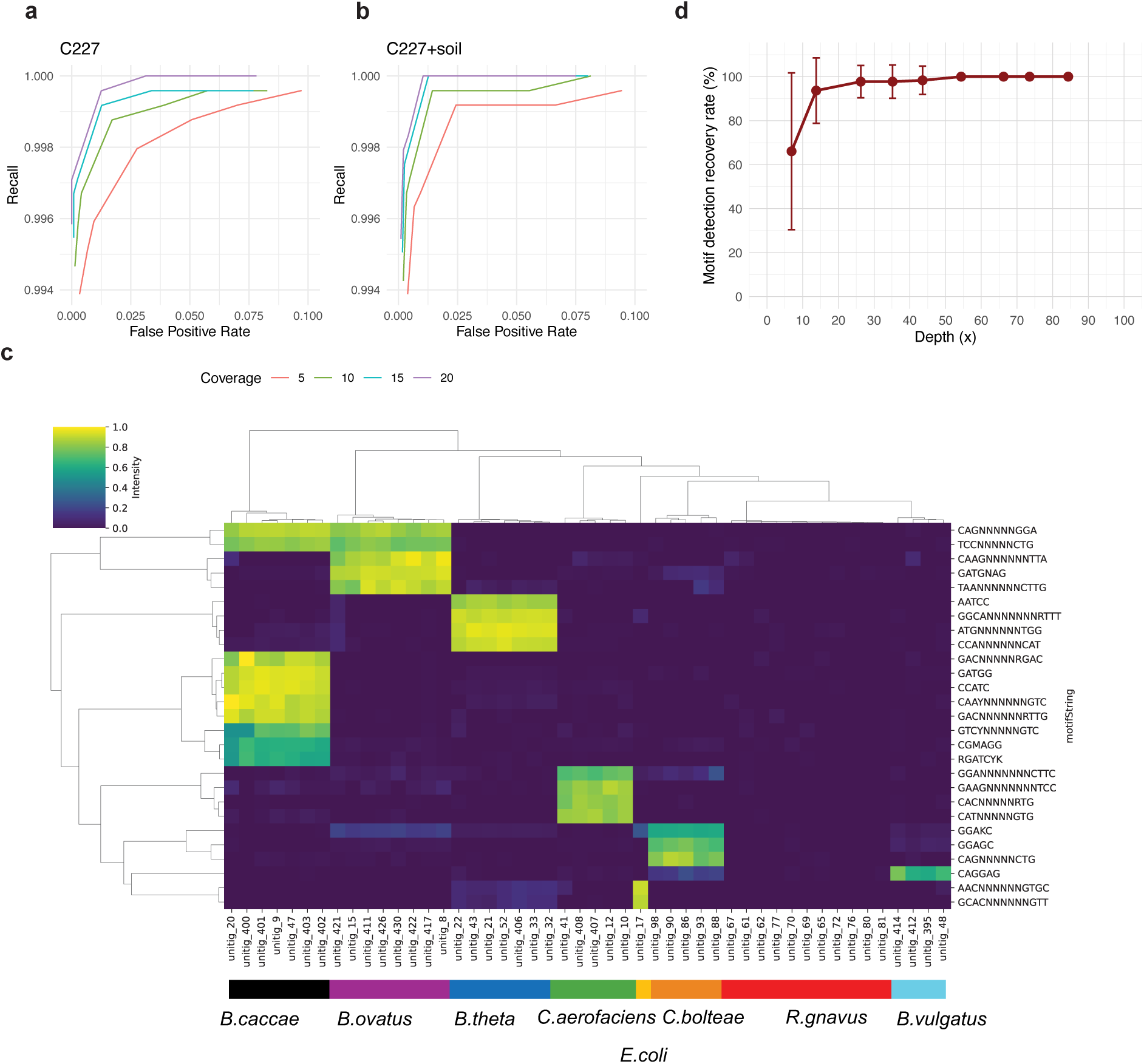
Benchmark of the DNA modification detection. **(a, b)** Recall and false positive rate of DNA base modification detection across a range of *P*-value cutoffs for both pure data **(a)** and data merged with soil samples **(b)**, where line colors distinguish different coverage cutoffs. **(c)** Modification profile of contigs within the JF8 mock community, with the bar colors below the heatmap indicating contigs belonging to different species and the heatmap color intensity representing the modification fraction for each specific motif. **(d)** Motif detection recovery rate across different depths in the PB24 dataset. Percentages of identified motifs compared to the total number of motifs detected with >50x sequencing depth. Error bar indicates standard deviations.

**Figure S2.**
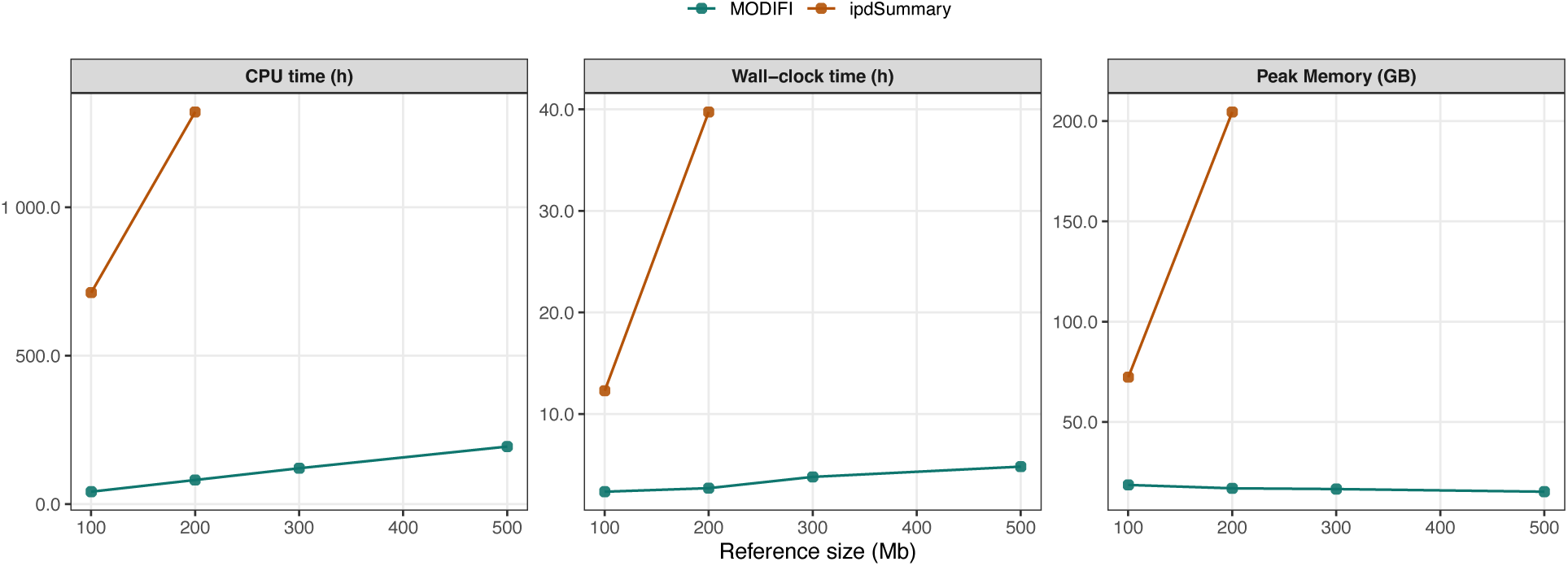
Computational resource consumption comparison between MODIFI and ipdSummary. The left, middle, and right panels exhibit CPU time, wall-clock time, and peak memory, respectively.

**Figure S3.**
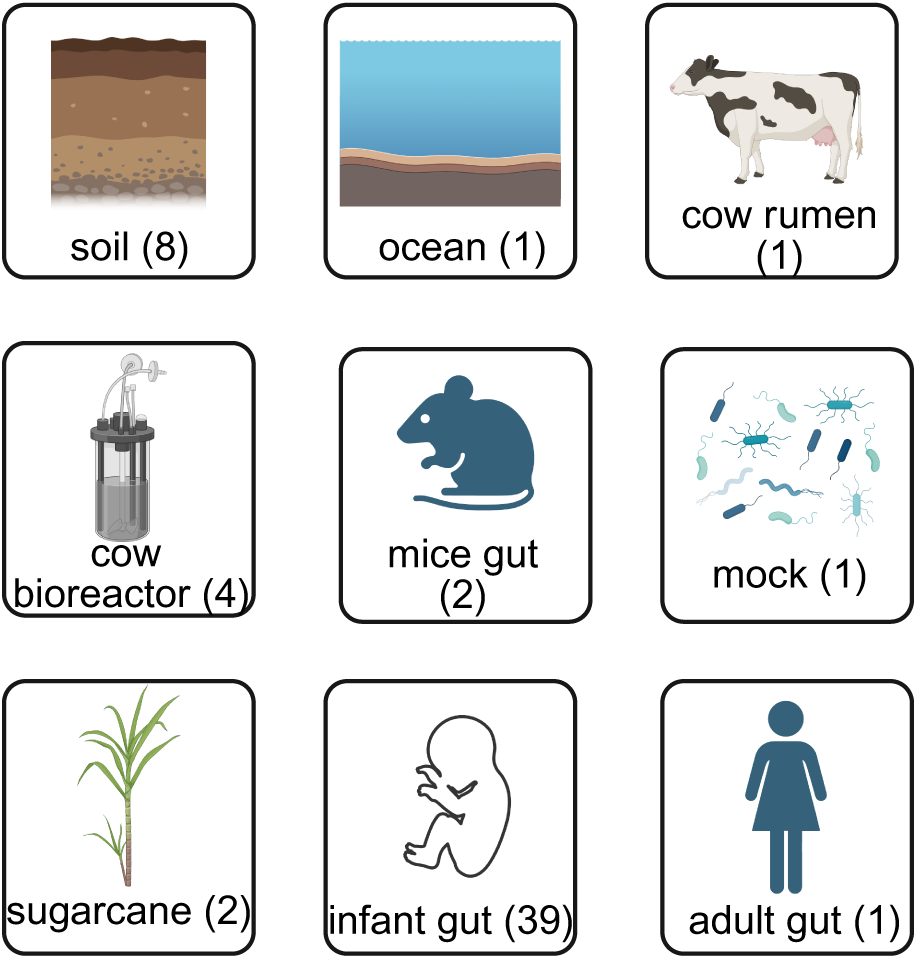
Distribution of metagenomic datasets across nine distinct habitats. Numerical values indicate the total number of samples processed for each habitat category.

**Figure S4.**
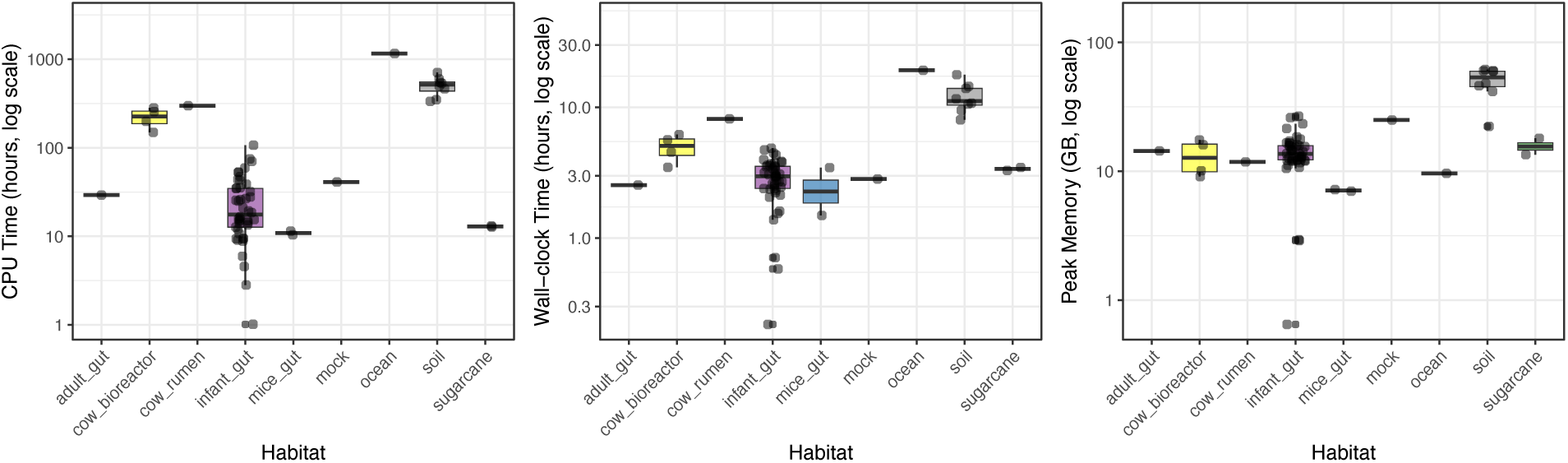
Computational resource consumption of MODIFI for DNA modification detection in 59 metagenomes across nine habitats. The left, middle, and right panels exhibit CPU time, wall-clock time, and peak memory, respectively. Colors distinguish different habitats.

**Figure S5.**
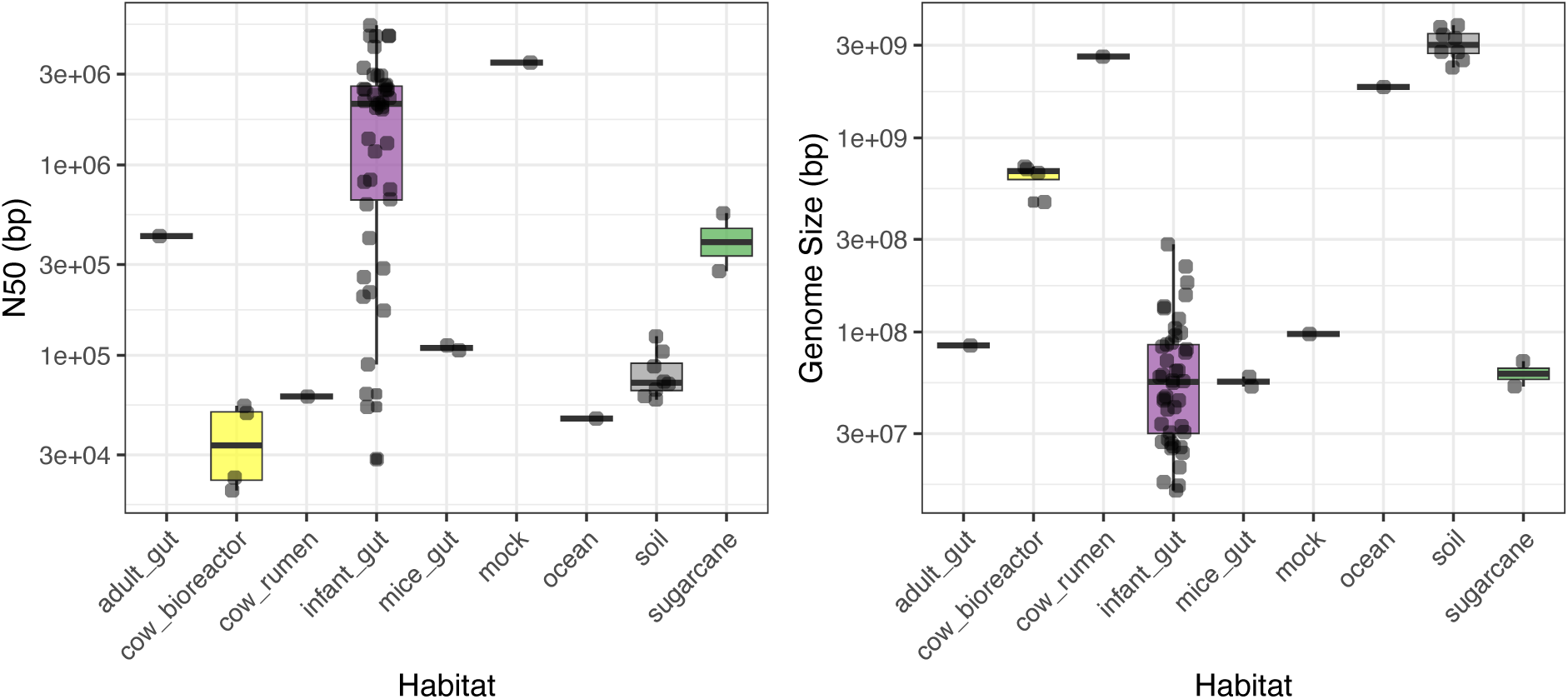
Boxplots of N50 (left) and metagenome size (right) of each sample across different habitats. The left panel represents the N50 values (bp) and the right panel represents the total metagenome size (bp) per sample.

**Figure S6.**
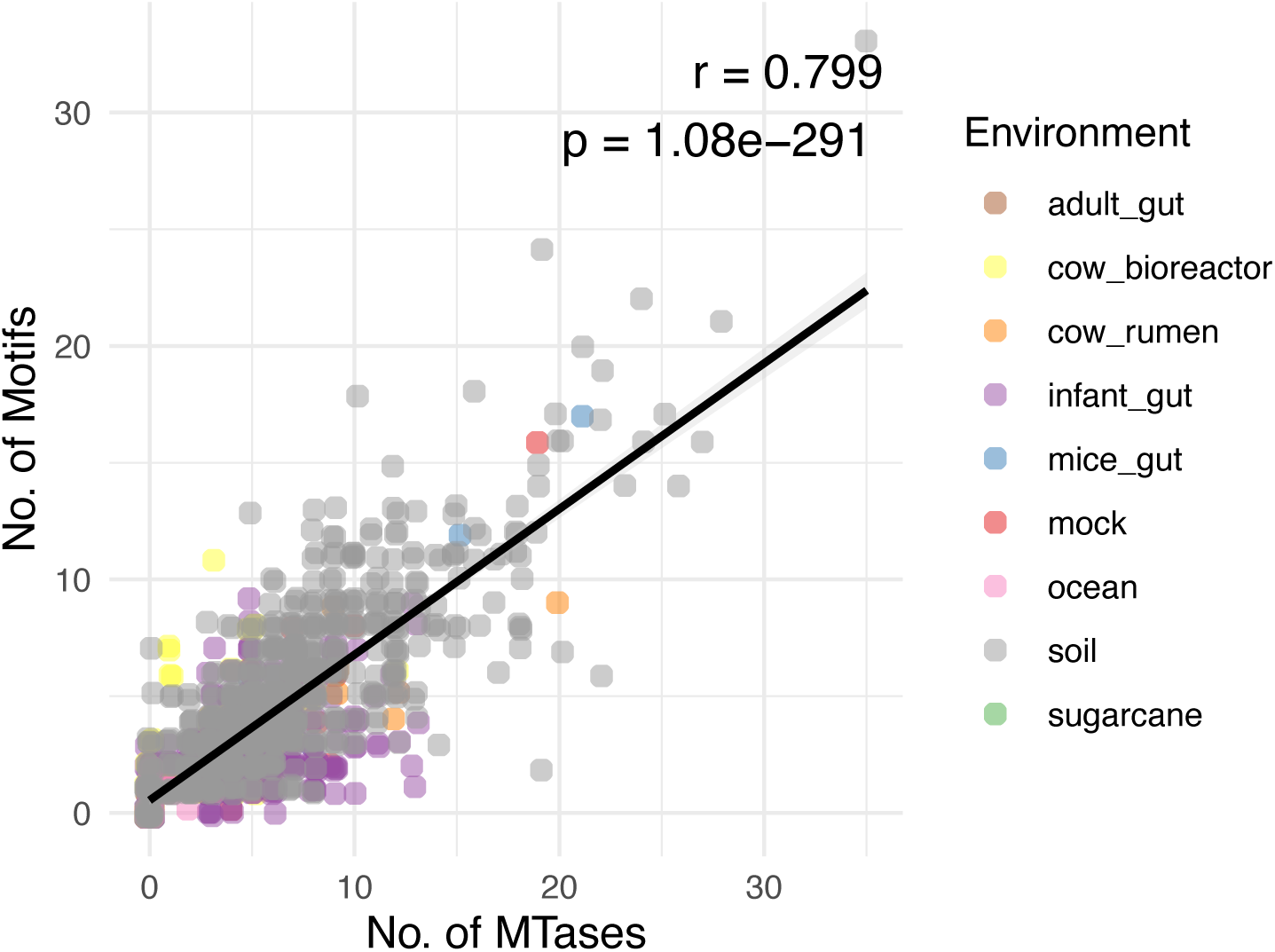
Correlation between MTase count and motif counts in each genome from metagenomes. Scatter plot showing the relationship between the per-genome motif number and MTase number across various metagenome samples. The solid black line represents the linear regression fit through the data points. The Pearson Correlation coefficient is displayed within the plot.

**Figure S7.**
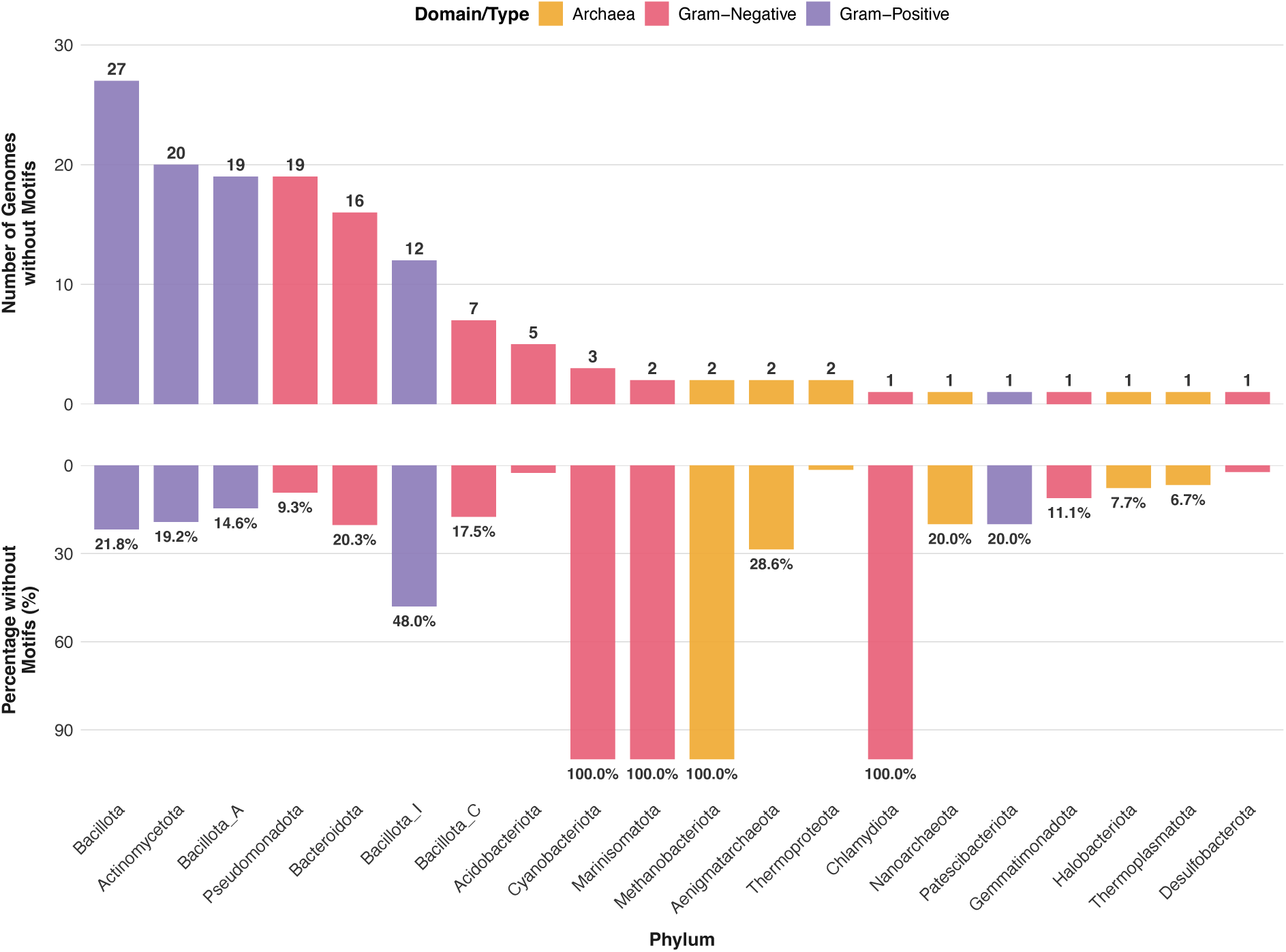
Distribution of unmodified genomes across phyla. The top bar plot displays the absolute number of unmodified genomes within each phylum, while the bottom bar plot indicates the corresponding proportion. Colors indicate the taxonomic classification: Gram-positive bacteria, Gram-negative bacteria, and Archaea.

**Figure S8.**
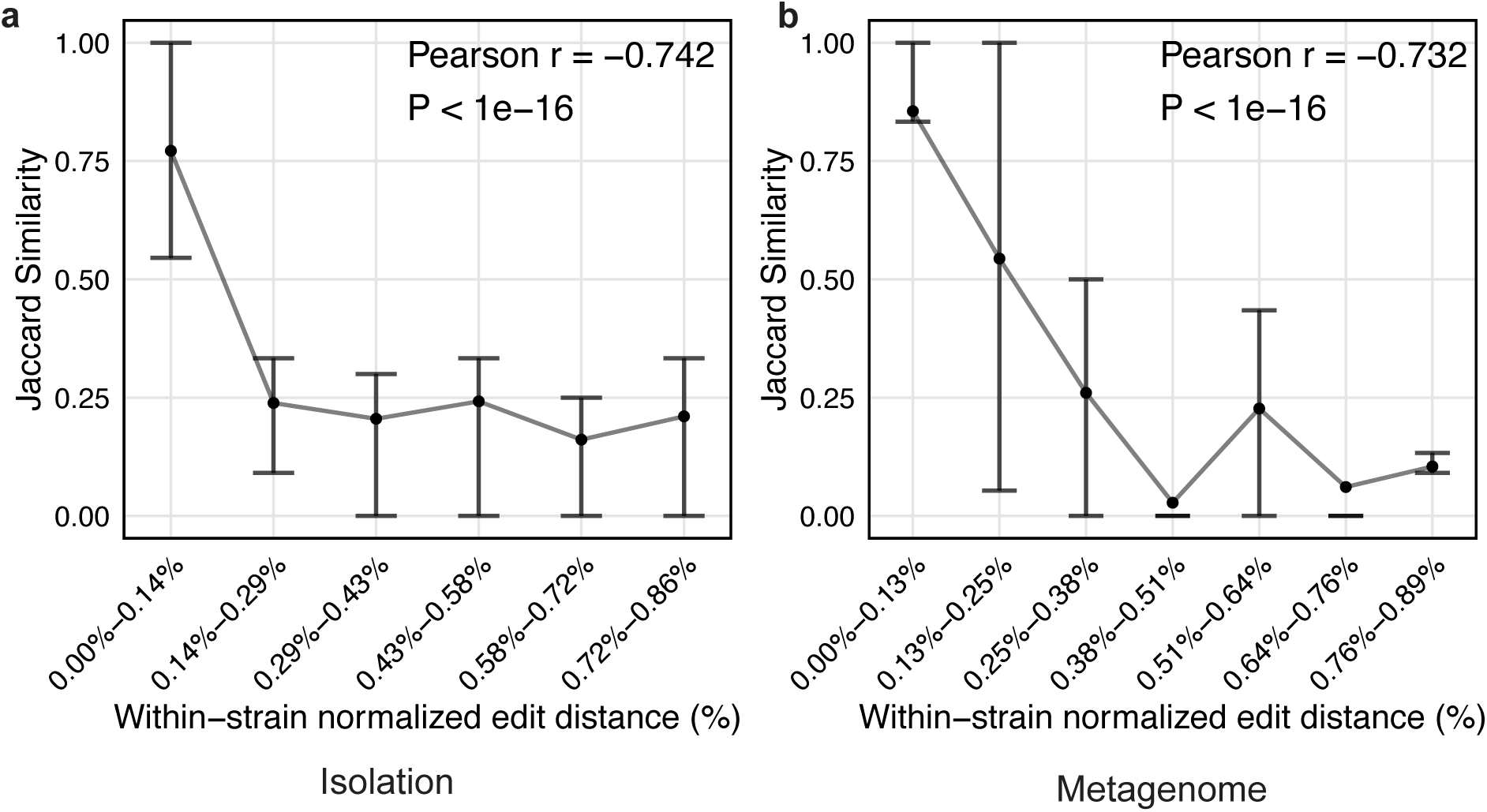
Correlation between modification similarity and genomic distance. Analysis of genome pairs within strains in isolation **(a)** and metagenomes **(b)**. The Jaccard similarity of modification motifs is plotted against the genome normalized edit distance for every genome pair, with normalized edit distance values grouped into discrete bins. Each point represents the mean Jaccard similarity for genome pairs within a normalized edit-distance bin, and vertical error bars indicate the interquartile range (25th-75th percentile) within that bin. The Pearson Correlation coefficient is displayed within the plot.

**Figure S9.**
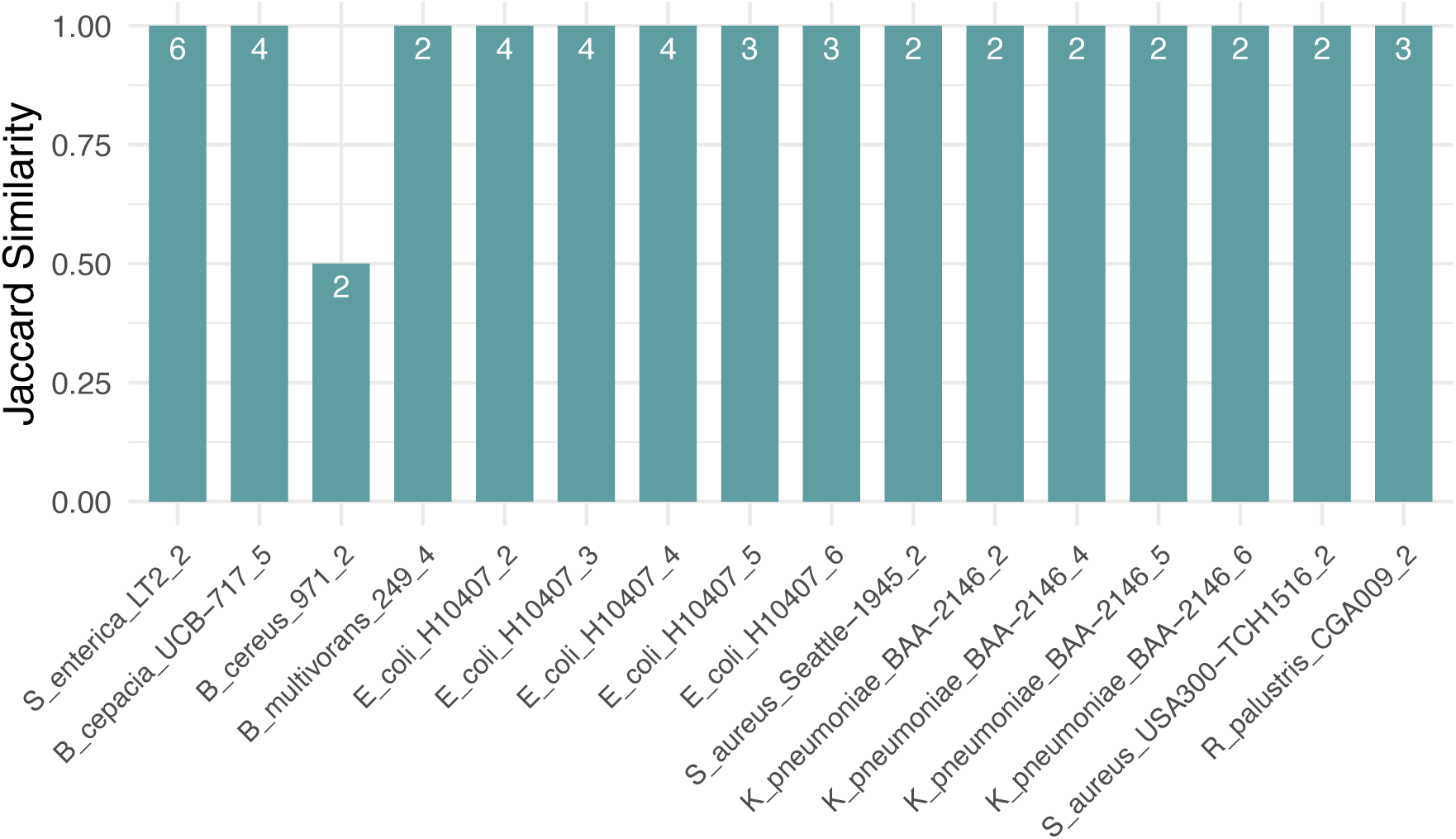
Modification similarity between ECE–host pairs in PB24 mock community. Jaccard similarity scores calculated for each plasmid and its corresponding host genome within the mock community; text labels within the bars specify the number of motifs used to calculate the similarity for each pair.

**Figure S10.**
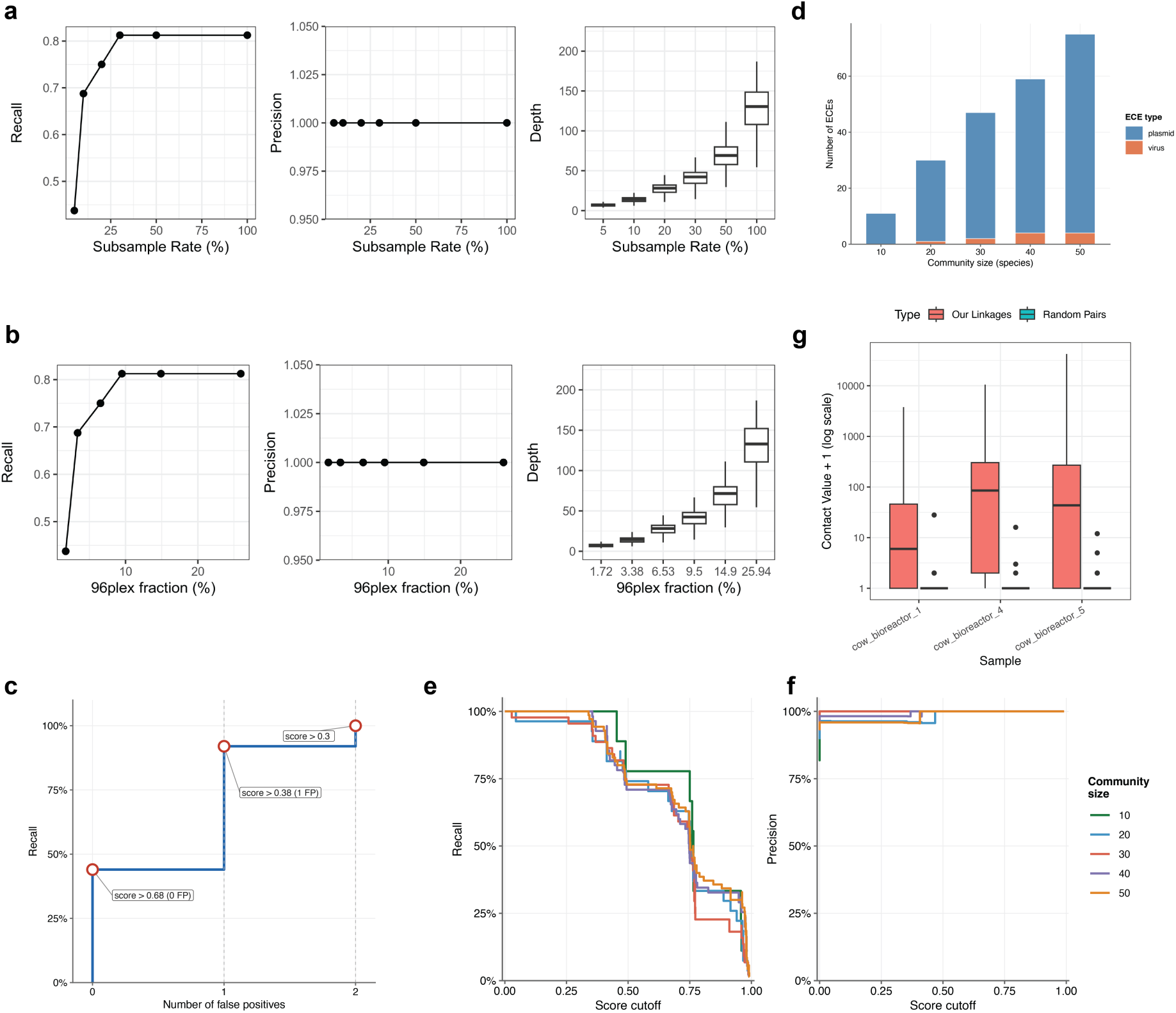
Performance evaluation of DNA modification-based ECE–host linkage prediction. **(a)** Benchmarking linkage accuracy using a PB24 mock community. Recall and precision of predicted host–ECE linkages are shown as a function of sequencing depth (subsample rate). **(b)** Benchmarking linkage accuracy in complex metagenomic contexts. Performance was evaluated by spiking PB24 mock community reads into an infant gut metagenome, with the 96plex fraction (x-axis) representing the proportion of mock community reads within the total merged dataset. **(c)** Evaluation of strain-resolved linkage inference, showing recall and the number of false-positive linkages across different cutoffs. **(d)** Number of ECEs included across different mock community sizes. **(e, f)** Recall **(e)** and precision **(f)** of ECE–host linkages using various linkage score cutoffs. Line colors distinguish different community sizes. **(g)** Boxplots of Hi-C contact values for motif-based ECE–host linkages and for random contig pairs. The plots illustrate the enrichment of Hi-C support for predicted pairs (red) compared to a null distribution of random expectations (teal). For visualization, contact values were log-transformed following the addition of a pseudocount 1 to each data point.

**Figure S11.**
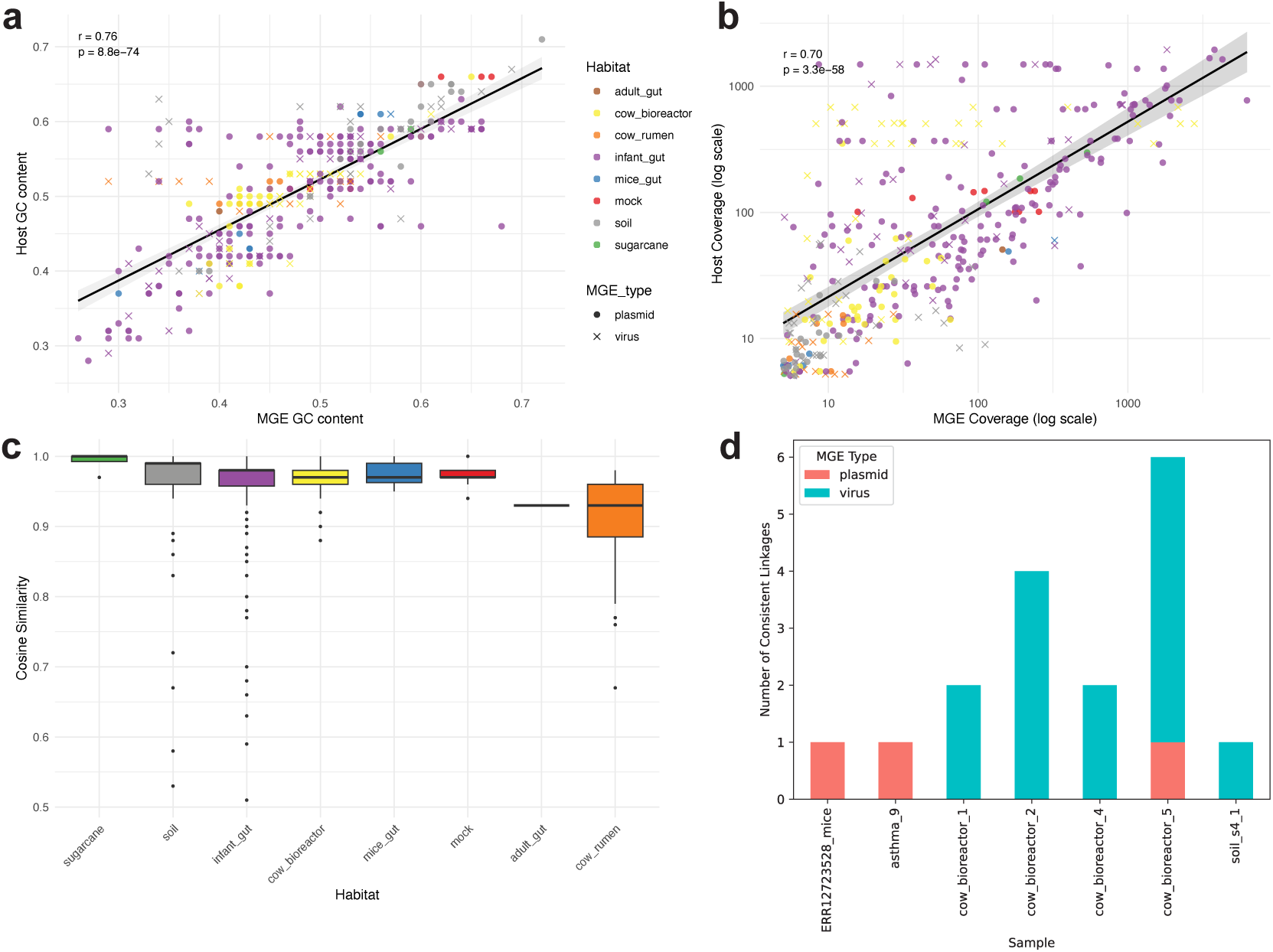
Characterization of ECEs and host linkages across diverse habitats. **(a)** Correlation of GC content between ECEs and hosts. Points are colored by habitat and shaped by ECE type (circles for plasmids, crosses for viruses). **(b)** Comparison of sequencing coverage (log scale) between ECEs and their hosts. **(c)** Boxplot of tetra-nucleotide similarity between ECE and host. **(d)** Validation of predicted linkages via CRISPR-Cas spacer matches. The bar chart displays the count of consistent supports for predicted linkages involving plasmids and viruses. Datasets with no consistent support were excluded from the analysis. Within **(a-b),** the Pearson Correlation coefficient is displayed, and black line represents the regression line.

**Figure S12.**
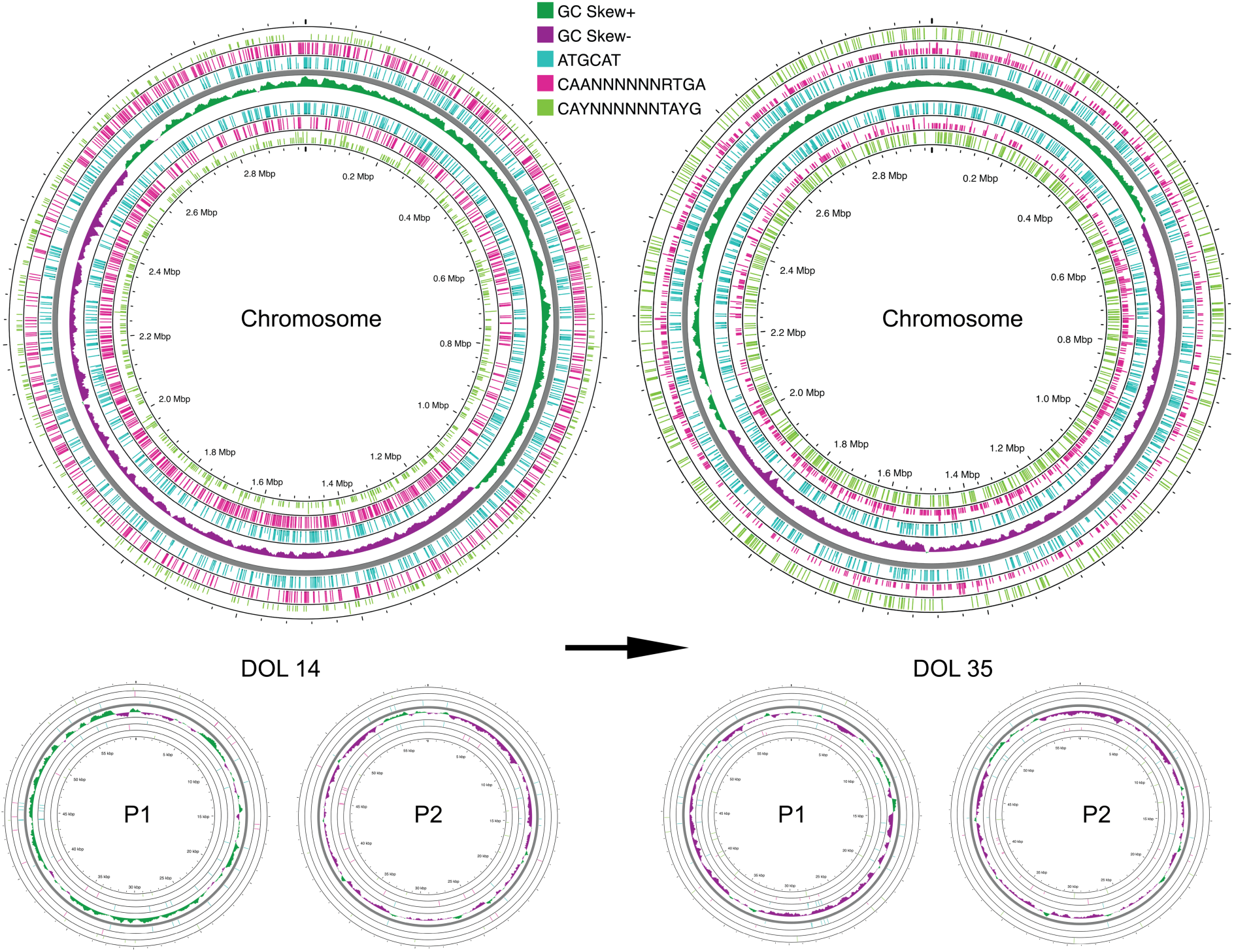
Illustration of modification shifts of *E. faecalis* and two associated plasmids at two time points. Each circos plot features a central track representing GC skew, flanked by modification motif sites for each individual DNA strand; at every motif occurrence site, full-length bars denote modified loci while half-length bars indicate unmodified loci, with bar colors utilized to distinguish between specific modification motifs.

**Figure S13.**
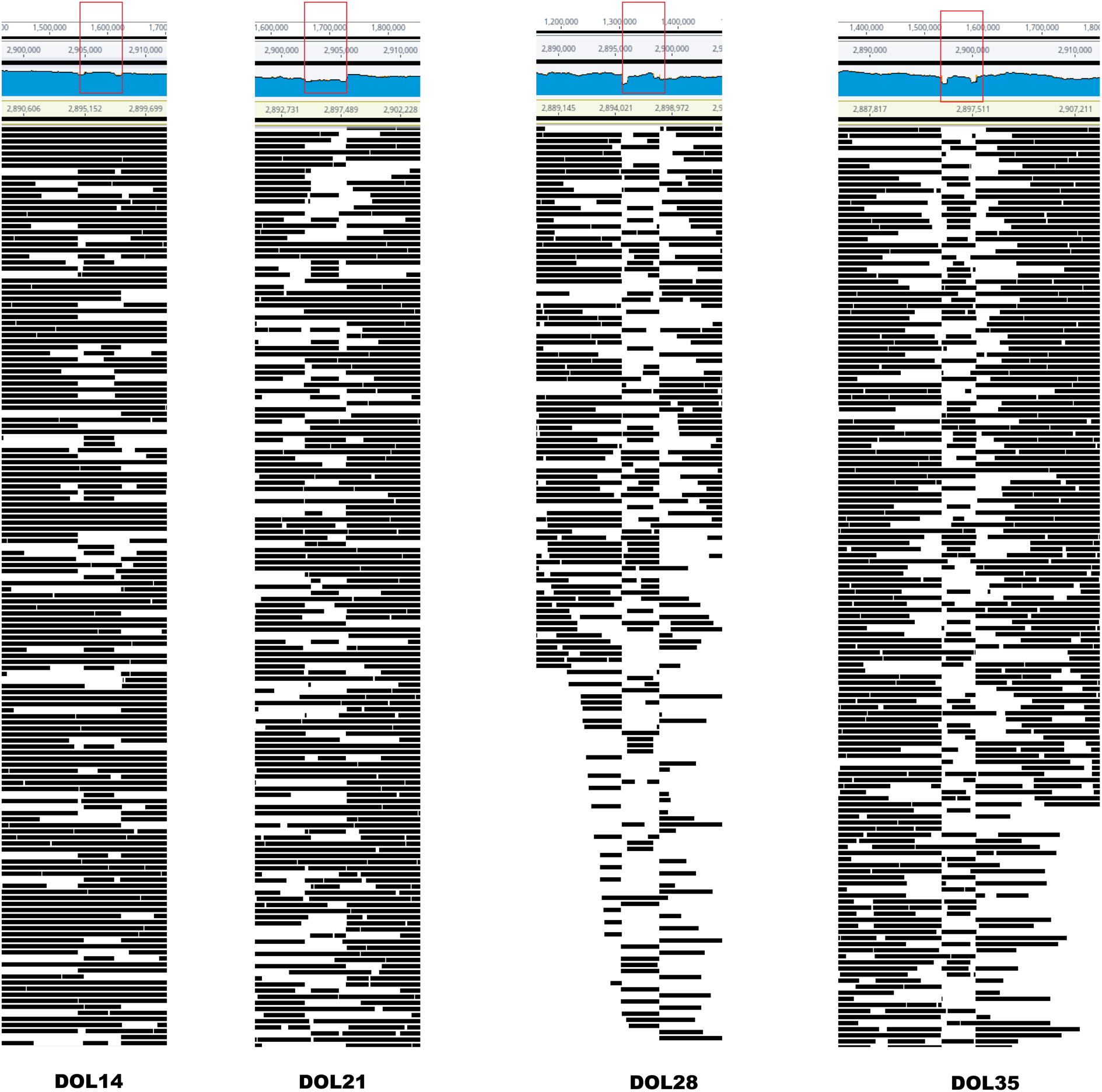
Temporal dynamics of read alignment to the *E. faecalis* inversion region. PacBio reads are mapped against the *E. faecalis* reference genome assembled at DOL14. Individual black bars represent single sequencing reads. The central region marked by the red rectangle represents the genomic inversion.

**Figure S14.**
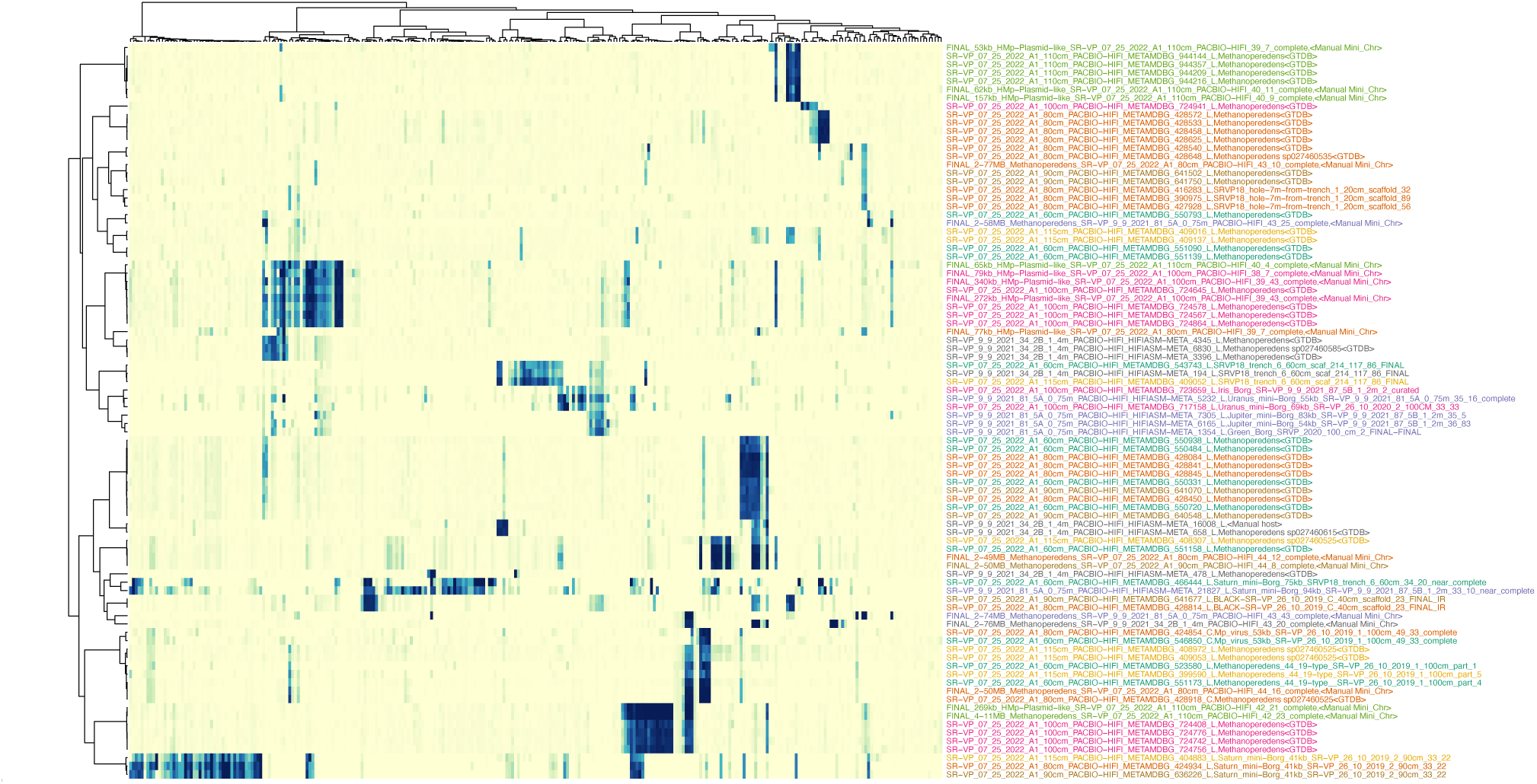
Heatmap of the DNA modifications across *Methanoperedens* and related ECEs. Each column is a distinct motif, and each row is a contig. The color gradient from yellow to blue represents an increasing modification fraction. Annotations on the right specify individual contig names and their corresponding genome classifications, with text colors utilized to differentiate between various source samples.

**Figure S15.**
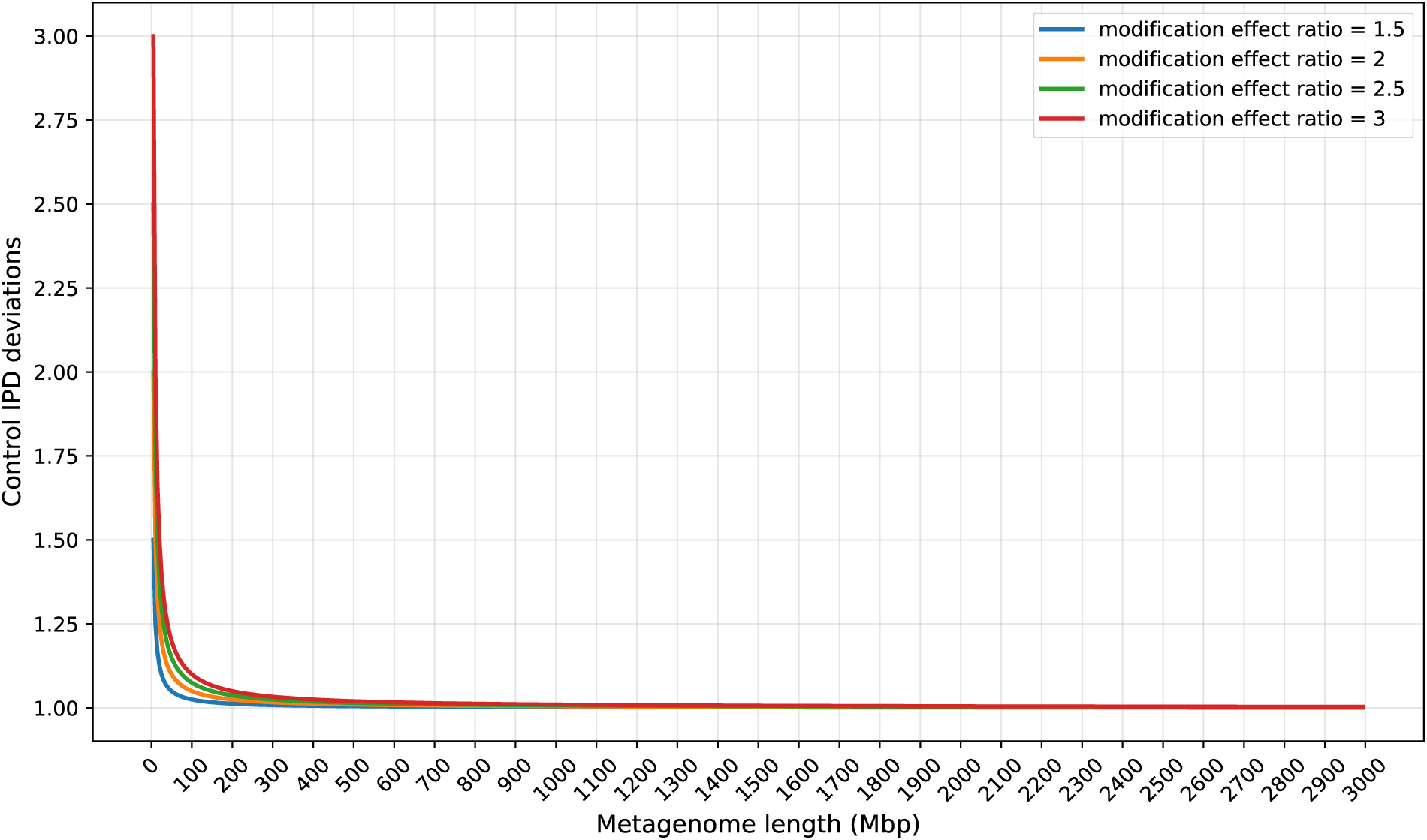
Evaluation of control IPD estimation accuracy across different metagenome lengths. Control IPD deviation indicates the estimated control IPD divided by real control IPD. The control IPD deviation converges toward 1 as metagenome length increases, indicating that the estimated control IPD effectively approaches the true unmodified baseline. Line colors indicate the modification effect ratio, defined as the ratio of modified to unmodified IPD.

## SUPPLEMENTARY TABLES

**Table S1 |** Motif detection in JF8 mock community

**Table S2 |** NCBI Accession numbers of isolates

**Table S3 |** Summary of environmental metagenomics and benchmark data

**Table S4 |** Summary of host genomes utilized for the removal of host-related reads prior to assembly

**Table S5 |** High-confident linkages from diverse environments

**Table S6 |** Distribution of motifs across different genomic regions in Borgs

**Table S7 |** Modification fractions across different genomic regions in Borgs

## References

1. Khedkar, S. et al. Landscape of mobile genetic elements and their antibiotic resistance cargo in prokaryotic genomes. Nucleic Acids Res. 50, 3155–3168 (2022).

2. Lang, A. S., Buchan, A. & Burrus, V. Interactions and evolutionary relationships among bacterial mobile genetic elements. Nat. Rev. Microbiol. 1–16 (2025).

3. Kang, D. D. et al. MetaBAT 2: an adaptive binning algorithm for robust and efficient genome reconstruction from metagenome assemblies. PeerJ 7, (2019).

4. Alneberg, J. et al. Binning metagenomic contigs by coverage and composition. Nat. Methods 11, 1144–1146 (2014).

5. Boeckaerts, D. et al. Prediction of Klebsiella phage-host specificity at the strain level. Nat. Commun. 15, 4355 (2024).

6. Tesson, F. et al. Systematic and quantitative view of the antiviral arsenal of prokaryotes. Nat. Commun. 13, 2561 (2022).

7. Roux, S. et al. Planetary-scale metagenomic search reveals new patterns of CRISPR targeting. bioRxiv 2025.06.12.659409 (2025) doi:10.1101/2025.06.12.659409.

8. Yang, Q. E., Gao, J. T., Zhou, S. G. & Walsh, T. R. Cutting-edge tools for unveiling the dynamics of plasmid-host interactions. Trends Microbiol. (2025) doi:10.1016/j.tim.2024.12.013.

9. Yu, F. B. et al. Microfluidic-based mini-metagenomics enables discovery of novel microbial lineages from complex environmental samples. Elife 6, (2017).

10. Beaulaurier, J., Schadt, E. E. & Fang, G. Deciphering bacterial epigenomes using modern sequencing technologies. Nat. Rev. Genet. 20, 157–172 (2019).

11. Beaulaurier, J. et al. Single molecule-level detection and long read-based phasing of epigenetic variations in bacterial methylomes. Nat. Commun. 6, 7438 (2015).

12. Oliveira, P. H. et al. Epigenomic characterization of Clostridioides difficile finds a conserved DNA methyltransferase that mediates sporulation and pathogenesis. Nat. Microbiol. 5, 166– 180 (2020).

13. Kobayashi, I., Nobusato, A., Kobayashi-Takahashi, N. & Uchiyama, I. Shaping the genome--restriction-modification systems as mobile genetic elements. Curr. Opin. Genet. Dev. 9, 649–656 (1999).

14. Beaulaurier, J. et al. Metagenomic binning and association of plasmids with bacterial host genomes using DNA methylation. Nat. Biotechnol. 36, 61–69 (2018).

15. Wilbanks, E. G. et al. Metagenomic methylation patterns resolve bacterial genomes of unusual size and structural complexity. ISME J. 16, 1921–1931 (2022).

16. Flusberg, B. A. et al. Direct detection of DNA methylation during single-molecule, real-time sequencing. Nat. Methods 7, 461–465 (2010).

17. Tourancheau, A., Mead, E. A., Zhang, X.-S. & Fang, G. Discovering multiple types of DNA methylation from bacteria and microbiome using nanopore sequencing. Nat. Methods 18, 491–498 (2021).

18. Clark, T. A. et al. Characterization of DNA methyltransferase specificities using single-molecule, real-time DNA sequencing. Nucleic Acids Res. 40, e29 (2012).

19. Trigodet, F., Sachdeva, R., Banfield, J. F. & Eren, A. M. Troubleshooting common errors in assemblies of long-read metagenomes. Nat. Biotechnol. 1–10 (2026).

20. Feng, Z. et al. Detecting DNA modifications from SMRT sequencing data by modeling sequence context dependence of polymerase kinetic. PLoS Comput. Biol. 9, e1002935 (2013).

21. Schadt, E. E. et al. Modeling kinetic rate variation in third generation DNA sequencing data to detect putative modifications to DNA bases. Genome Res. 23, 129–141 (2013).

22. Jha, A. et al. DNA-m6A calling and integrated long-read epigenetic and genetic analysis with fibertools. Genome Res. 34, 1976–1986 (2024).

23. Li, H., Niu, J., Sheng, Y., Liu, Y. & Gao, S. SMAC: identifying DNA N^6^-methyladenine (6mA) at the single-molecule level using SMRT CCS data. bioRxiv (2024) doi:10.1101/2024.11.13.623492.

24. Seong, H. J., Roux, S., Hwang, C. Y. & Sul, W. J. Marine DNA methylation patterns are associated with microbial community composition and inform virus-host dynamics. Microbiome 10, 157 (2022).

25. Ni, M., et al. Epigenetic phase variation in the gut microbiome enhances bacterial adaptation. bioRxivorg (2025) doi:10.1101/2025.01.11.632565.

26. Blow, M. J. et al. The epigenomic landscape of prokaryotes. PLoS Genet. 12, e1005854 (2016).

27. Roberts, R. J., Vincze, T., Posfai, J. & Macelis, D. REBASE--a database for DNA restriction and modification: enzymes, genes and genomes. Nucleic Acids Res. 43, D298–9 (2015).

28. Al-Shayeb, B. et al. Borgs are giant genetic elements with potential to expand metabolic capacity. Nature 610, 731–736 (2022).

29. Schoelmerich, M. C. et al. Borg extrachromosomal elements of methane-oxidizing archaea have conserved and expressed genetic repertoires. Nat. Commun. 15, 5414 (2024).

30. Banfield, J. F. et al. Convergent evolution of viral-like Borg archaeal extrachromosomal elements and giant eukaryotic viruses. Nat. Commun. 16, 10641 (2025).

31. Shi, L.-D. et al. Jumbo circular extrachromosomal elements of methane-oxidizing archaea with variably extensive metabolic and defense gene repertoires. bioRxiv 2026.01.21.700959 Preprint at 10.64898/2026.01.21.700959 (2026).

32. Schoelmerich, M. C., Sachdeva, R., West-Roberts, J., Waldburger, L. & Banfield, J. F. Tandem repeats in giant archaeal Borg elements undergo rapid evolution and create new intrinsically disordered regions in proteins. PLoS Biol. 21, e3001980 (2023).

33. Murray, I. A. et al. The methylomes of six bacteria. Nucleic Acids Res. 40, 11450–11462 (2012).

34. Hiraoka, S. et al. Diverse DNA modification in marine prokaryotic and viral communities. Nucleic Acids Res. 50, 1531–1550 (2022).

35. Albers, S.-V. & Meyer, B. H. The archaeal cell envelope. Nat. Rev. Microbiol. 9, 414–426 (2011).

36. Giovannoni, S. J. et al. Genome streamlining in a cosmopolitan oceanic bacterium. Science 309, 1242–1245 (2005).

37. Wigington, C. H. et al. Re-examination of the relationship between marine virus and microbial cell abundances. Nat. Microbiol. 1, 15024 (2016).

38. Yan, M., Banfield, J. & Sachdeva, R. Revealing the pervasive landscape of MGE-host interactions in situ with single-cell genomics. bioRxiv 2025.12. 20.690322 (2025) doi:10.64898/2025.12.20.690322.

39. Meaden, S., Westra, E. R. & Fineran, P. C. Phage defence-system abundances vary across environments and increase with viral density. Philos. Trans. R. Soc. Lond. B Biol. Sci. 380, 20240069 (2025).

40. Rodriguez-Valera, F. et al. Explaining microbial population genomics through phage predation. Nat. Rev. Microbiol. 7, 828–836 (2009).

41. Rocha, E. P. C. & Bikard, D. Microbial defenses against mobile genetic elements and viruses: Who defends whom from what? PLoS Biol. 20, e3001514 (2022).

42. Gorzynski, J. et al. Bacterial defense systems and host ecology drive the evolution of intra-species lineages. Cell Rep. 45, 116957 (2026).

43. Tisza, M. J. et al. Roving methyltransferases generate a mosaic epigenetic landscape and influence evolution in Bacteroides fragilis group. Nat. Commun. 14, 4082 (2023).

44. Chanin, R. B. et al. Intragenic DNA inversions expand bacterial coding capacity. Nature 634, 234–242 (2024).

45. Gorrie, C. L. et al. Genomic dissection of Klebsiella pneumoniae infections in hospital patients reveals insights into an opportunistic pathogen. Nat. Commun. 13, 3017 (2022).

46. Gelsinger, D. R. et al. Metagenomic editing of commensal bacteria in vivo using CRISPR-associated transposases. Science 390, eadx7604 (2025).

47. Rubin, B. E. et al. Species- and site-specific genome editing in complex bacterial communities. Nat. Microbiol. 7, 34–47 (2022).

48. Kungulovski, G. & Jeltsch, A. Epigenome editing: State of the art, concepts, and perspectives. Trends Genet. 32, 101–113 (2016).

49. Takahashi, M. et al. Host-encoded DNA methyltransferases modify the epigenome and host tropism of invading phages. iScience 28, 112264 (2025).

50. SMRT Link. PacBio https://www.pacb.com/smrt-link/ (2023).

51. Wang, S., Jiang, Y., Che, L., Wang, R. H. & Li, S. C. Enhancing insights into diseases through horizontal gene transfer event detection from gut microbiome. Nucleic Acids Res. 52, e61 (2024).

52. Li, T., Zhang, X., Luo, F., Wu, F.-X. & Wang, J. MultiMotifMaker: A multi-thread tool for identifying DNA methylation motifs from PacBio reads. IEEE/ACM Trans. Comput. Biol. Bioinform. 17, 220–225 (2020).

53. Tejada, J. N. et al. Prevention and cure of murine C. difficile infection by a Lachnospiraceae strain. Gut Microbes 16, 2392872 (2024).

54. Lin, H. et al. Metagenome-based diversity and functional analysis of culturable microbes in sugarcane. Microbiol. Spectr. 13, e0198224 (2025).

55. Guitor, A. K. et al. Megaplasmids associate with Escherichia coli and other Enterobacteriaceae. bioRxivorg 2025.09.30.679422 (2025) doi:10.1101/2025.09.30.679422.

56. Morowitz, M. J. et al. The NICU Antibiotics and Outcomes (NANO) trial: a randomized multicenter clinical trial assessing empiric antibiotics and clinical outcomes in newborn preterm infants. Trials 23, 428 (2022).

57. Cabana, M. D., McKean, M., Wong, A. R., Chao, C. & Caughey, A. B. Examining the hygiene hypothesis: the Trial of Infant Probiotic Supplementation. Paediatr. Perinat. Epidemiol. 21 **Suppl 3**, 23–28 (2007).

58. Fujimura, K. E. et al. House dust exposure mediates gut microbiome Lactobacillus enrichment and airway immune defense against allergens and virus infection. Proc. Natl. Acad. Sci. U. S. A. 111, 805–810 (2014).

59. Li, H. Minimap2: pairwise alignment for nucleotide sequences. Bioinformatics 34, 3094– 3100 (2018).

60. Feng, X., Cheng, H., Portik, D. & Li, H. Metagenome assembly of high-fidelity long reads with hifiasm-meta. Nat. Methods 19, 671–674 (2022).

61. Cheng, H., Concepcion, G. T., Feng, X., Zhang, H. & Li, H. Haplotype-resolved de novo assembly using phased assembly graphs with hifiasm. Nat. Methods 18, 170–175 (2021).

62. Chklovski, A., Parks, D. H., Woodcroft, B. J. & Tyson, G. W. CheckM2: a rapid, scalable and accurate tool for assessing microbial genome quality using machine learning. Nat. Methods 20, 1203–1212 (2023).

63. Chaumeil, P.-A., Mussig, A. J., Hugenholtz, P. & Parks, D. H. GTDB-Tk: a toolkit to classify genomes with the Genome Taxonomy Database. Bioinformatics 36, 1925–1927 (2019).

64. Letunic, I. & Bork, P. Interactive Tree of Life (iTOL) v6: recent updates to the phylogenetic tree display and annotation tool. Nucleic Acids Res. 52, W78–W82 (2024).

65. Grant, J. R. et al. Proksee: in-depth characterization and visualization of bacterial genomes. Nucleic Acids Res. 51, W484–W492 (2023).

66. Camargo, A. P. et al. Identification of mobile genetic elements with geNomad. Nat. Biotechnol. 42, 1303–1312 (2024).

67. Olm, M. R., Brown, C. T., Brooks, B. & Banfield, J. F. dRep: a tool for fast and accurate genomic comparisons that enables improved genome recovery from metagenomes through de-replication. ISME J. 11, 2864–2868 (2017).

68. Kurtz, S. et al. Versatile and open software for comparing large genomes. Genome Biol. 5, R12 (2004).

69. DeMaere, M. Z. & Darling, A. E. bin3C: exploiting Hi-C sequencing data to accurately resolve metagenome-assembled genomes. Genome Biol. 20, 46 (2019).

70. Bland, C. et al. CRISPR recognition tool (CRT): a tool for automatic detection of clustered regularly interspaced palindromic repeats. BMC Bioinformatics 8, 209 (2007).

71. Altschul, S. F. et al. Gapped BLAST and PSI-BLAST: a new generation of protein database search programs. Nucleic Acids Res. 25, 3389–3402 (1997).

72. Crits-Christoph, A., Kang, S. C., Lee, H. H. & Ostrov, N. MicrobeMod: A computational toolkit for identifying prokaryotic methylation and restriction-modification with nanopore sequencing. bioRxiv (2023) doi:10.1101/2023.11.13.566931.

73. Bastian, M., Heymann, S. & Jacomy, M. Gephi: An open source software for exploring and manipulating networks. Proceedings of the International AAAI Conference on Web and Social Media 3, 361–362 (2009).

74. Hyatt, D. et al. Prodigal: prokaryotic gene recognition and translation initiation site identification. BMC Bioinformatics 11, 119 (2010).

